# Meta-omics profiling of full-scale groundwater rapid sand filters explains stratification of iron, ammonium and manganese removals

**DOI:** 10.1101/2022.12.07.519464

**Authors:** Francesc Corbera-Rubio, Michele Laureni, Nienke Koudijs, Simon Müller, Theo van Alen, Frank Schoonenberg, Sebastian Lücker, Martin Pabst, Mark C.M. van Loosdrecht, Doris van Halem

## Abstract

Rapid sand filters (RSF) are an established and widely applied technology for groundwater treatment. Yet, the underlying interwoven biological and physical-chemical reactions controlling the sequential removal of iron, ammonia and manganese remain poorly understood. To resolve the contribution and interactions between the individual reactions, we studied two full-scale drinking water treatment plant configurations, namely (*i*) one dual-media (anthracite and quartz sand) filter and (*ii*) two single-media (quartz sand) filters in series. *In situ* and *ex situ* activity tests were combined with mineral coating characterization and metagenome-guided metaproteomics along the depth of each filter. Both plants exhibited comparable performances and process compartmentalization, with most of ammonium and manganese removal occurring only after complete iron depletion. Within each compartment, the homogeneity of the media coating and genome-based microbial composition highlighted the effect of backwashing on filter media mixing. In stark contrast, intra-compartment contaminant removal was highly stratified following decreasing substrate availability along the filter height. This apparent and long-standing conflict was resolved by quantifying the expressed proteome at different filter heights, revealing a consistent stratification of proteins catalysing ammonia oxidation and protein-based relative abundances of nitrifying genera. This implies that microorganisms adapt their protein pool to the available nutrient load at a faster rate than the backwash mixing frequency. Ultimately, these results show the unique and complementary potential of metaproteomics to understand metabolic adaptations and interactions in highly dynamic ecosystems.

## 1. Introduction

Anaerobic groundwater is an excellent drinking water source due to its microbiological safety, stable temperature and composition (Giordano, 2009; Katsanou & Karapanagioti, 2019; de Vet, 2011). Rapid sand filters (RSFs), preceded by an aeration step, are the most commonly applied technology for groundwater treatment. In RSFs, the main groundwater contaminants – soluble iron (Fe^2+^), ammonium (NH_4_^+^) and manganese (Mn^2+^) - are sequentially removed in a series of interwoven biological and physical-chemical reactions (Bourgine et al., 1994). While the latter have been the subject of decades of research, our understanding of the involved microbiology is limited. Our current understanding is that Fe^2+^ is removed both chemically and biologically (Van Beek et al., 2012), Mn^2+^ via chemical autocatalytic oxidation and to some extent biologically (Breda, Søborg, et al., 2019), and NH_4_^+^exclusively biologically (Tekerlekopoulou et al., 2013). However, not only the impact of microbial activity on the removal processes and their stratification along the height of RSFs is largely unknown, but also the individual physiologies of the involved microorganisms remain elusive (Gülay et al., 2016). Deepening our knowledge of the underlying microbiology is paramount to improve current RSF operation and to design and implement novel, resource-efficient systems.

The first characterization of nitrifying biomass stratification in RSFs dates back to the beginning of this century (Kihn et al., 2000). In their pioneering work, Kihn and colleagues observed an uneven distribution of nitrifying activity along a full-scale RSF using *ex-situ* batch incubations. Subsequently, molecular methods, such as fluorescence *in situ* hybridization (Lydmark et al., 2006), 16S rRNA gene cloning and sequencing (Qin et al., 2007), and denaturing gradient gel electrophoresis (de Vet et al., 2009) have been used to investigate the diversity of nitrifying bacteria and to estimate the relative abundance of ammonia-oxidizing bacteria and archaea. Furthermore, quantitative polymerase chain reaction (qPCR) has been applied to determine the distribution of nitrifying and denitrifying microorganisms in RSFs (Bai et al., 2013), linking absolute microbial abundances to *in situ* ammonium removal capacity (Tatari et al., 2016; Tatari et al., 2017). Recently, shotgun metagenomics has established itself as the predominant approach for the characterization of the composition and functional potential of complex microbiomes, and led to remarkable discoveries in RSFs such as the complete oxidation of ammonia by a single organism (comammox) (Palomo et al., 2016; Pinto et al., 2016) or the metabolic-coupling between methane- and ammonia-oxidizing communities (Poghosyan et al. (2020)). On the contrary, information about biological iron and manganese oxidation in RSFs is scarce, primarily due to our limited understanding of the underlying metabolic pathways (Ilbert & Bonnefoy, 2013; Romano et al., 2017). In spite of this, the presence and importance of iron- and manganese-oxidizing organisms in RSFs has repeatedly been shown by culture-dependent (Burger et al., 2008; Nitzsche et al., 2015; S. Qin et al., 2009; L. Yang et al., 2014) and culture-independent studies (Bruins et al., 2017; Gülay et al., 2018; Marcus et al., 2017; H. Yang et al., 2020), *ex situ* batch incubations (Breda, Ramsay, et al., 2019), and metagenomics (Hu et al., 2020; Palomo et al., 2016). However, mechanistic insights linking filter performance with actual microbial activities have not been established to date.

Recent technical advancements made the characterization of the entire pool of proteins expressed in complex communities feasible (Van Den Bossche et al., 2021; Wilmes & Bond, 2004). Proteins are the biological entities catalyzing metabolic reactions. Thus, beyond genome-based approaches, their identification and quantification allow for the simultaneous resolution of the identity and actual metabolic function of core community members (Kleiner, 2019). Protein abundances also provide a more accurate estimation of the relative biomass contribution of different populations in a community (Kleikamp et al., 2021; Kleiner et al., 2017). Over the past decades, metaproteomics has been successfully applied to characterize microbiomes from different environments, including marine (Stokke et al., 2012), soil (Schulze et al., 2005), wastewater (Kleikamp et al., 2022) and freshwater (Hanson & Madsen, 2015) ecosystems. Moreover, differential protein expression has been used to characterize microbiomes’ physiological responses to rapid environmental changes (Wilmes et al., 2015), making metaproteomics particularly relevant to study dynamic environments such as RSF systems.

RSFs are regularly backwashed to remove particles that have been trapped during operation. As a result, biomass-colonized filter media grains are displaced along the filter height and exposed to different substrate loads at every cycle (Ramsay et al., 2021). Consistently, metagenome-based studies commonly report even distributions of core taxa along filter heights (Bai et al., 2013; Tatari et al., 2017), and conventional RSFs models consider them as homogeneous systems (Uhl & Gimbel, 2000). In contrast, and irrespective of the backwash, contaminant removal stratification along the filter is commonly reported (Gude et al., 2016; Tatari et al., 2016). Within this framework, we hypothesize that genetically homogeneous RSFs’ microbial communities hold the ability to rapidly modify their pool of expressed proteins to metabolically respond to the environmental conditions encountered at a given filter height. To this end, we quantified the *in-situ* removal of iron, manganese, and ammonium, and resolved the taxonomy and function of the microbial communities along the filter height of two full-scale RSF plants, one dual-media (anthracite and sand) single filter system and one comprising two single-media sand filters in series.

## 2. Materials and methods

### 2.1. Sample collection

Filter media and water samples were collected from two groundwater-fed drinking water treatment plants (DWTP) operated by the company Vitens. The first one, located in Druten, was a single filter (SF) with a dual media bed of 0.6 m anthracite on top and 2 m of quartz sand at the bottom. The second one, located in Hasselo, was a double filter (DF) consisting of two single-media 2 m high quartz sand filters (DF1 and DF2) connected in series. The filter media was sampled with a peat sampler at three different heights in each filter (approximately every 0.5 m), referred to as top (t), middle (m) and bottom (b) sections (Figure 1). Sampling from the bottom section of DF1 (*i.e*., DF1_b) was technically not possible. Filter media samples were taken at the end of the operational cycle before backwashing. Influent, effluent, and filter-bed porewater samples were immediately filtered (0.45 µm), stored at 4 °C, and measured within 12 h. Samples for total iron and manganese quantification were acidified to pH < 1 with 69 % ultrapure nitric acid. pH, O_2_ concentration, temperature and redox potential were measured on-site using a multimeter (Multi 3630 IDS, Xylem Analytics, Germany). Operational parameters and raw water characteristics for each filter are shown in Table 1.

**Table 1.**
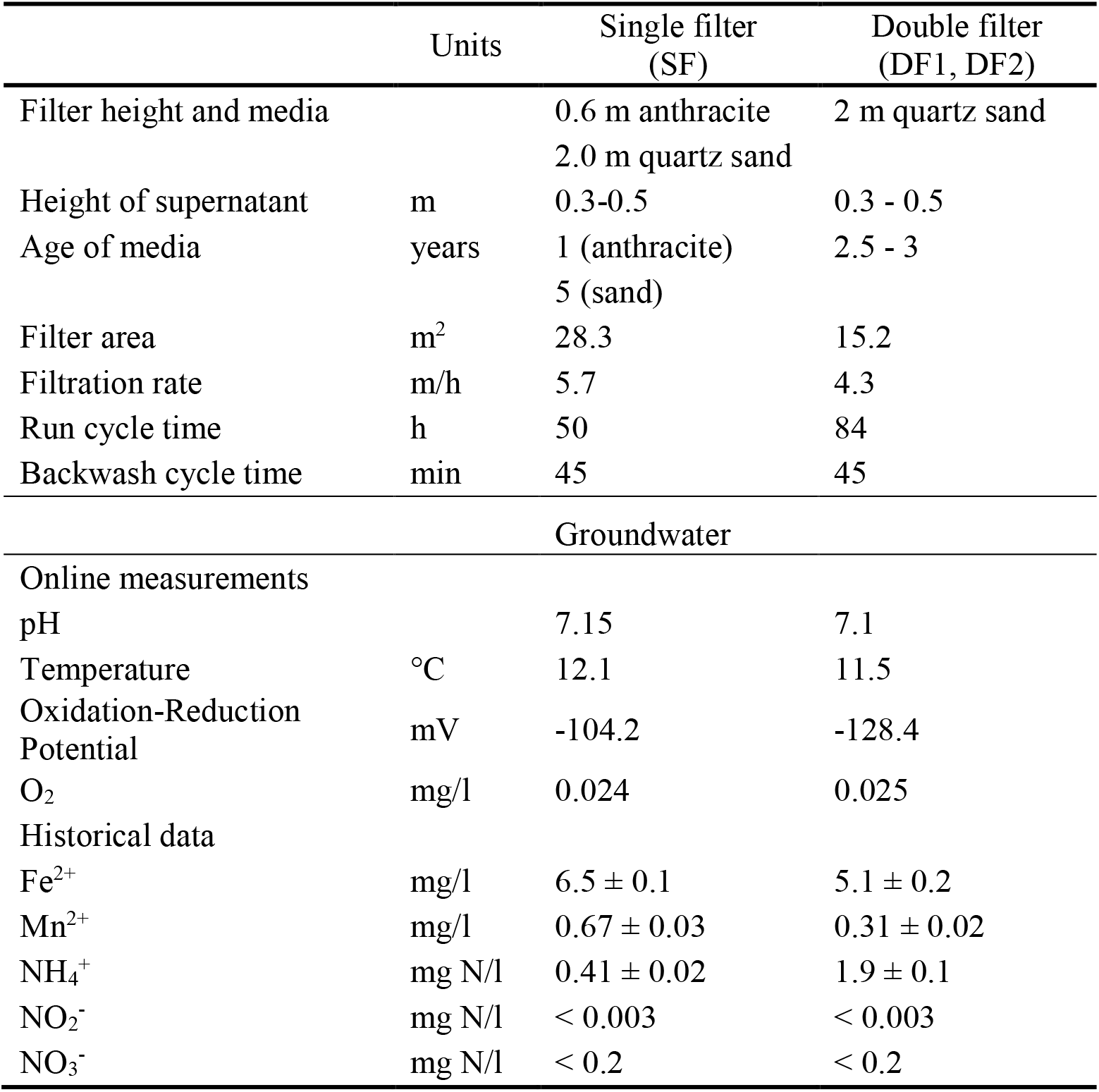
Operational parameters of the DWTPs and the characteristics of the respective groundwaters. Historical data refer to average and standard deviation of the weekly concentrations between November 2018 and July 2020. For the double filter system, data refer to the influent to the first filter.

|  | Units | Single filter (SF) | Double filter (DF1, DF2) |
| --- | --- | --- | --- |
| Filter height and media |  | 0.6 m anthracite<br>2.0 m quartz sand | 2 m quartz sand |
| Height of supernatant | m | 0.3-0.5 | 0.3 - 0.5 |
| Age of media | years | 1 (anthracite)<br>5 (sand) | 2.5 - 3 |
| Filter area | m<sup>2</sup> | 28.3 | 15.2 |
| Filtration rate | m/h | 5.7 | 4.3 |
| Run cycle time | h | 50 | 84 |
| Backwash cycle time | min | 45 | 45 |
| Groundwater |  |  |  |
| Online measurements |  |  |  |
| pH |  | 7.15 | 7.1 |
| Temperature | °C | 12.1 | 11.5 |
| Oxidation-Reduction Potential | mV | -104.2 | -128.4 |
| O<sub>2</sub> | mg/l | 0.024 | 0.025 |
| Historical data |  |  |  |
| Fe<sup>2+</sup> | mg/l | 6.5 ± 0.1 | 5.1 ± 0.2 |
| Mn<sup>2+</sup> | mg/l | 0.67 ± 0.03 | 0.31 ± 0.02 |
| NH<sub>4</sub><sup>+</sup> | mg N/l | 0.41 ± 0.02 | 1.9 ± 0.1 |
| NO<sub>2</sub><sup>-</sup> | mg N/l | < 0.003 | < 0.003 |
| NO<sub>3</sub><sup>-</sup> | mg N/l | < 0.2 | < 0.2 |

**Figure 1.**
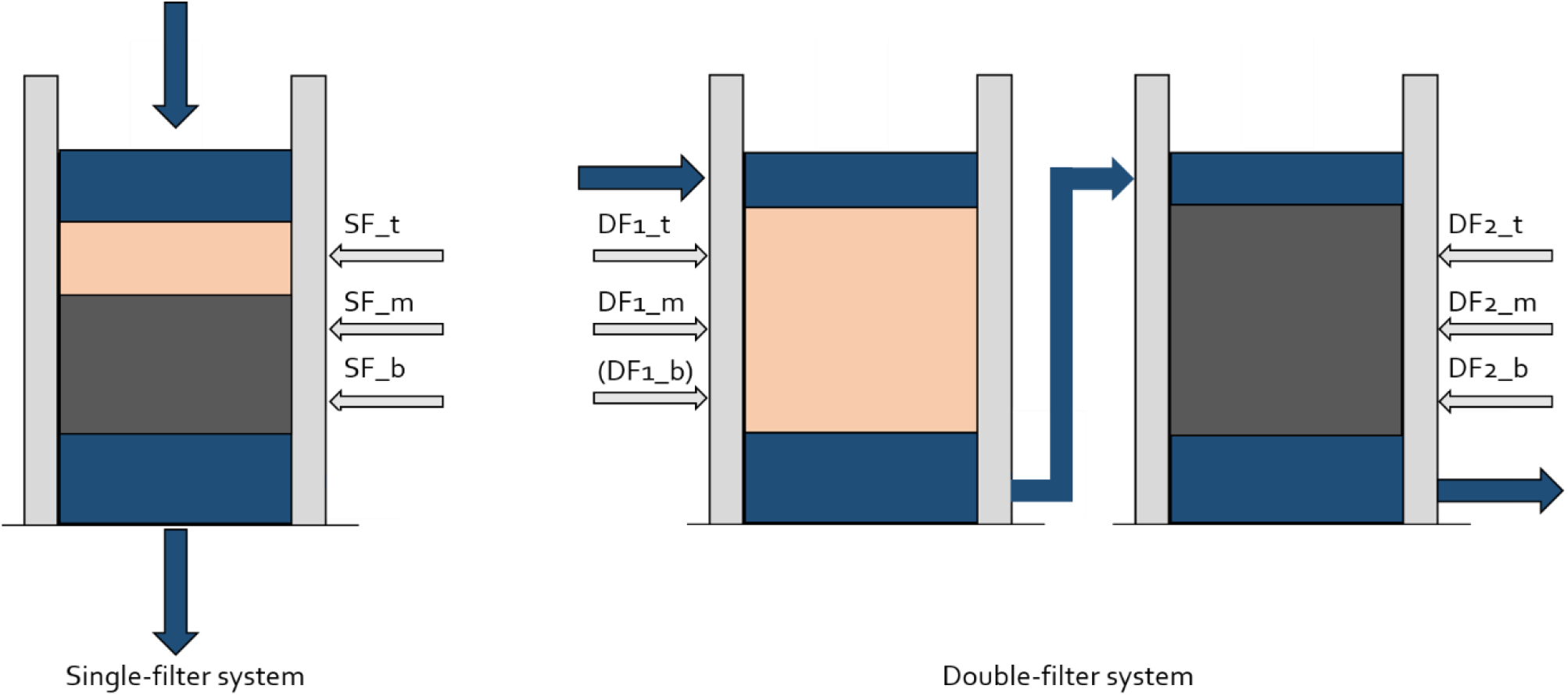
Schematic illustration of the two drinking water treatment plants. Each individual filter is characterized at three different heights: top (t), middle (m) and bottom (b). Colors of the filter media material reproduce the color of the coating (beige and dark grey for iron and manganese oxides, respectively), and reflect the compartmentalization of the removal processes.

### 2.2. Ex situ *ammonium and manganese maximum removal rates*

The maximum ammonium and manganese removal rates of the filter media were determined in batch. 4 g of wet filter media, 200 mL of tap water, and 100 μl of trace element solution (L^-1^; 15g EDTA, 4.5g ZnSO_4_ ·7H_2_O, 4.5g CaCl_2_·2H_2_O, 3g FeSO_4_·7H_2_O, 1g H_3_BO_3_, 0.84g MnCl_2_·2H_2_O, 0.3g CoCl_2_·6H_2_O, 0.3g CuSO_4_·5H_2_O, 0.4g Na_2_MoO_4_·2H_2_O, 0.1g KI) were mixed in 300 mL shake flasks. After an acclimatization period of 30 min at 25 °C and 150 rpm, each flask was spiked with 3 ml of 100 mg NH_4_^+^-N L^-1^ NH_4_Cl or 100 mg Mn^2+^ L^-1^ MnCl_2_·4H_2_O (Sigma Aldrich, Saint Louis, Missouri USA). Liquid samples (1 ml) were taken at different time intervals throughout the entire process. Maximum removal rates per mass (wet weight) of filter media were calculated by interpolation of the concentration profiles and converted into volumetric rates using the measured filter media densities (Table S4). Biological ammonia oxidation was suppressed by overnight incubation at 50 °C with 0.05 g penicillin g L^-1^ (Sigma Aldrich) (H. Yang et al., 2020) and confirmed with ATP measurements (see below).

### 2.3. Water quality analyses

Samples for ammonium, nitrite, and nitrate quantification were immediately filtered through a 0.2 µm nanopore filter and measured within 12 h using photometric analysis (Gallery Discrete Analyzer, Thermo Fischer Scientific, Waltham, Massachusetts, USA). Samples for dissolved iron and manganese quantification were immediately filtered through a 0.2 μm nanopore filter and analysed within 12 h. Raw water samples were filtered after a minimum of 16 hours of acidification for total iron and manganese quantification. Iron and manganese were quantified by ICP-MS (Analytik Jena, Jena, Germany).

For ATP detection, 1 ml miliQ H_2_O was added to 1 g wet weight of sand. The samples were sonicated for 1 min at an output power of 15 W and a frequency of 20 kHz (ultrasonic homogenizer, Qsonica sonicators, Newtown, Connecticut, USA). Samples were subsequently filtered through a 0.22 μm nanopore filter. The concentration of ATP in the liquid was determined in duplicates using an ATP analyzer as described by the manufacturer (Clean-Trace™ Luminometer NG3, 3M, Maplewood, Minnesota USA).

### 2.4. Filter-bed coating characterisation and visualization

Coating extraction was carried out in triplicate as described elsewhere (Claff et al., 2010) with minor modifications. 1.5 g of wet filter media were frozen overnight at -80 °C and freeze-dried for 48 h (Alpha 1-4 LD plus, Christ, Osterode am Harz, Germany). Samples were immersed in 40 ml citrate buffered dithionite solution for 4 h, centrifuged at 4000 rpm for 10 minutes, and filtered through a 0.45 µm pore size filter. Iron was measured using the 3500-Fe B: Phenanthroline method (APHA, 2018), and manganese colorimetrically (LCK 532, Hach Lange, Tiel, The Netherlands). Light microscopy images were taken with a VHX-5000 digital microscope (Keyence, Osaka, Japan) with an VH-Z20UR lens.

### 2.5. DNA extraction

Nucleic acid extraction was carried out using the MagMAX CORE Nucleic Acid Purification Kit (Applied Biosystems, Thermo Fisher Scientific). A pre-treatment step using bead beating tubes from MagMAX CORE Mechanical Lysis Module was introduced to improve DNA recovery. In duplicates, 0.25 g of sample, 350 µL lysis solution, and 10 µL proteinase K were mixed and bead-beaten for 2 × 30 seconds (Bead Mill Homogenizer, BioSpec, Bartlesville, Oklahoma, USA), and then centrifuged for 2-7 minutes at 10000 × *g* until the supernatant was clear. Next, the supernatants of two tubes were combined in a clean tube, mixed with 450 µL Bead/PK mix and vortexed for 10 min. The tubes were placed on a magnetic stand for 1 min and the supernatant was removed. The samples were washed twice by adding 500 µL of ‘wash solution 1/2’, and vortexed for 1 minute prior to supernatant removal on the magnetic stand. After the second washing step, samples were air-dried for 5 minutes. 90 µL elution buffer were added and samples were vortexed for 10 min. After 2 min on the magnetic stand, the supernatants were transferred to clean tubes. Afterwards, all samples were purified using the GeneJET PCR Purification kit following the manufacturer’s protocol (Thermo Scientific, Thermo Fisher Scientific). DNA was quantified with a Qubit 4 Fluorometer and Qubit dsDNA HS assay kit (Invitrogen, Thermo Fisher Scientific).

### 2.6. Library preparation and sequencing

Metagenomic library preparation was performed using the Nextera XT kit (Illumina, San Diego, California U.S.A.) according to the manufacturer’s instructions. Enzymatic tagmentation was performed starting with 1 ng of DNA, followed by incorporation of the indexed adapters and PCR amplification. After purification of the amplified library using AMPure XP beads (Beckman Coulter, Indianapolis, USA), the libraries were checked for quality and size distribution using an Agilent 2100 Bioanalyzer and the High sensitivity DNA kit (Agilent, San Diego, USA). Quantitation of the library was performed by Qubit using the Qubit dsDNA HS Assay Kit (Invitrogen, Thermo Fisher Scientific, USA). The libraries were pooled, denatured, and sequenced with Illumina MiSeq (San Diego, California USA). Paired end sequencing of 2 × 300 base pairs was performed using the MiSeq Reagent Kit v3 (San Diego, California USA) according to the manufacturer’s instructions.

### 2.7. Assembly, functional annotation, and taxonomic classification

Raw sequencing data was quality-filtered and trimmed using Trimmomatic v0.39 (HEADCROP:16 LEADING:3 TRAILING:5 SLIDINGWINDOW:4:10 CROP:240 MINLEN:35) (Bolger et al., 2014). Sequencing data quality was analyzed using FastQC v0.11.7 before and after trimming (Andrews, 2010). MEGAHIT v1.2.9 was used to assemble the quality-filtered and trimmed reads into contigs with default settings (Li et al., 2015). Raw reads were mapped back to the assembled contigs using BWA-MEM2 (Vasimuddin et al., 2019). SAMtools v1.14 was used to determine contig coverage and for indexing with default settings (Li et al., 2009). Taxonomic classification of each contig was performed using the Contig Annotation Tool (CAT) with default settings and the NCBI database (Von Meijenfeldt et al., 2019a). Ultimately, the relative abundance of each taxa was calculated by matching the CAT results and the read mapping information (SAMtools), expressed as percentage of the sum of the trimmed reads mapping to the contigs of each taxa compared to the total number of trimmed reads (Albertsen et al., 2013). Functional annotation was done with GhostKoala v2.2 (Kanehisa et al., 2016) against the Kyota Enciclopedia of Genes and Genomes (KEGG; accessed September 2021). FeGenie (Garber et al., 2020) was used to refine the annotation of genes involved in iron metabolism using the metagenomics (‘-meta’) settings. Genes encoding the three subunits of the particulate methane monooxygenase (*pmoABC*) and ammonia monooxygenase (*amoABC*) were translated into protein sequences and differentiated by constructing phylogenetic trees in MEGA11 using *MUSCLE* as alignment tool and the Neighbor-Joining method with the *WAG* substitution model (Hall, 2013) (Figure S2, Figure S3, Figure S4). Reference sequences were extracted from the National Center for Biotechnology Information database (2021_09), ensuring a wide representation of all beta- and gammaproteobacterial ammonia-oxidizing bacteria (Purkhold et al., 2000), and methanotrophs affiliated with the Alpha- and Gammaproteobacteria and *Verrumicrobia* (Kalyuzhnaya et al., 2019) in the resulting phylogenetic trees. Nitrite oxidoreductases (*nxrAB*) and nitrate reductases (*narGH*) were manually identified based on taxonomy after their protein sequences were blasted using ‘blastp’ against the SwissProt database (v. 2022_02). In analogy, the nitrite reductases *nirK* and *nirS* were taxonomically assigned to nitrifying or denitrifying organisms. Twelve published Mn(II)-oxidizing genes (Hu et al., 2020) were used as reference database for manganese oxidation. Protein assemblies of each sample were aligned to the database using local blastp v2.13 with e-value < 1e-6, percentage identity > 35 % (Rost, 1999) and coverage > 70 % (Garber et al., 2020). RStudio v1.4.1106 was used for data analysis and visualization (RStudio Team, 2021). Raw DNA sequences can be found on the National Center for Biotechnology Information (NCBI) website under BioProject PRJNA830362.

### 2.8. Protein extraction, proteolytic digestion, and shotgun proteomic analysis

For protein extraction, 150 mg of filter (sand) material was mixed with 125 µL of B-PER reagent (78243, Thermo Scientific) and 125 µL 1 M TEAB buffer (50 mM TEAB, 1 % w/w NaDOC, adjusted to pH 8.0). The mixture was briefly vortexed and exposed to shaking using a Mini Bead Beater 16 (BioSpec Products) for 3 min. Afterwards, the sample was exposed to one freeze/thaw cycle at -80 °C and +80 °C for 15 and 5 min, respectively. The samples were centrifuged for 10 min at 10000 rpm at 4 °C, and the supernatant was transferred into a clean LoBind tube Eppendorf tube (Eppendorf, Hamburg, Germany). Another 125 µL of B-PER reagent and 125 µL 1 M TEAB buffer was added to the sand sample, vortexed and sonicated (Branson 5510, sonication mode, at room temperature) for 5 min. The sample was then centrifuged for 10 min at 10000 rpm at 4 °C, and the supernatant was collected and pooled with the first supernatant. Extracted proteins were precipitated by adding 200 µL ice-cold acetone to the supernatant, vortexing and incubation at -20 °C for 1 h. The protein pellet was collected by centrifugation at 14000 rpm at 4 °C for 15 min. The supernatant was carefully removed from the pellet using a pipette. The pellet was dissolved in 200 mM ammonium bicarbonate containing 6 M urea. Disulfide bonds were reduced using 10 mM DTT (dithiothreitol) and sulfhydryl groups were blocked using 20 mM IAA (iodoacetamide). The solution was diluted to below 1 M urea and digested using sequencing grade Trypsin (Promega). The proteolytic peptides were finally desalted using an Oasis HLB solid phase extraction well plate (Waters) according to the manufacturer’s protocol. An aliquot corresponding to approximately 250 ng of protein digest was analyzed with a shotgun proteomics approach as described earlier (Kleikamp et al., 2021). Briefly, the peptides were analyzed using a nano-liquid-chromatography system consisting of an EASY nano-LC 1200, equipped with an Acclaim PepMap RSLC RP C18 separation column (50 um × 150 mm, 2 µm, 100 Å) and a QE plus Orbitrap mass spectrometer (Thermo Fisher, Germany). The flow rate was maintained at 350 nL over a linear gradient from 5 % to 30 % solvent B over 65 minutes and finally to 60 % B over 20 min. Solvent A was ultrapure H_2_O containing 0.1 % v/v formic acid, and solvent B consisted of 80 % acetonitrile in H_2_O and 0.1 % v/v formic acid. The Orbitrap was operated in data-dependent acquisition (DDA) mode acquiring peptide signals from 385-1250 m/z at 70K resolution. The top 10 signals were isolated at a window of 2.0 m/z and fragmented at a NCE of 28 at 17.5K resolution with an AGC target of 2e5 and a max IT 75 ms. The collected mass spectrometric raw data were analyzed combined against the metagenomics reference sequence database built from the assembled contigs using PEAKS Studio 10 (Bioinformatics Solutions, Canada) in a two-round search approach. The first-round database search using the clustered database (892436 proteins) was used to construct a focused protein sequence database, considering one missed cleavage, and carbamidomethylation as fixed modification, and allowing 15 ppm parent ion and 0.02 Da fragment ion mass error. The focused database (containing 9729 proteins) was then used in a second-round search, considering 3 missed cleavages, carbamidomethylation as fixed and methionine oxidation and N/Q deamidation as variable modifications. Peptide spectrum matches were filtered for 5 % false discovery rate (FDR) and protein identifications with at least 2 unique peptides were considered as significant. Functional annotations (KO identifiers) and taxonomic lineages were obtained by GhostKoala (https://www.kegg.jp/ghostkoala) and by Diamond (Buchfink et al., 2014) using the NCBI nr protein sequence reference database (Index of /blast/db/FASTA (nih.gov)). For Diamond annotations, the lineage of the lowest common ancestor was determined from the top 10 sequence alignments.

Comparison of the relative abundance of proteins between samples was performed using “relative spectral counts” (*i.e*., normalized spectral counts, defined as spectral counts per protein divided by the molecular weight, divided by the sum of spectral counts of all proteins of all samples). Since protein extraction was performed from filter media of equal mass (150 mg per sample), the protein abundance of sample SF_t was multiplied upfront by 0.76 to correct for the density difference between anthracite (1.3 g/cm^3^) and quartz sand (1.7 g/cm^3^). For sake of simplicity, further corrections for differences in nutrient loads between systems, resulting from different superficial velocities and filter volumes (Table 1), were omitted as they did not yield significantly different results. To identify the functional guilds of interest, all proteins were taxonomically classified using their best hit in the NCBI and KEGG database and subsequently grouped at genus level, and the resulting genera were manually classified based on their most probable energy source, iron, nitrogen or “others”, based on the literature (Table S3). Similarly, the contribution of a specific protein was estimated by summing up the relative abundances of all proteins with the same annotation.

## 3. Results

### 3.1. In situ *oxidation of Fe^2+^, NH_4_^+^, Mn^2+^ display similar compartmentalisation in both configurations*

The concentration profiles of ammonium, iron and manganese along the studied treatment processes are shown in Figure 1A and 2B. Iron was completely removed (< 0.04 mg Fe^2+^/L) in the top layer of the single-filter system (*i.e*., anthracite layer, SF_t) and in the first filter of the double-filter system (DF1). Consistently, manganese consumption only started after iron depletion in both systems, namely in SF_m and DF2_t, and was complete (< 0.002 mg Mn^2+^/L) at the end of each treatment process. In analogy, most ammonium was removed after iron depletion, with < 30% removal in SF_t and DF1.

The observed process compartmentalization was also reflected in each filter section in the total mass of iron and manganese oxides, the end products of Fe^2+^ and Mn^2+^ oxidation (Van Beek et al., 2012) (Figure 2C, D). The coating in SF_t and DF1 mainly consisted of iron (118 ± 8 mg_Fe_/g_media_), while the coating in SF_m, SF_b, and DF2 mainly consisted of manganese (33 ± 6 mg_Mn_/g_media_). In SF, anthracite and quartz sand remained clearly separated after over one year of operation despite frequent backwashing cycles. Nevertheless, trace amounts of anthracite particles were found in the lower filter sections and might explain the slightly higher iron coating content in SF_m and SF_b as compared to DF2 (Figure 2C, D).

**Figure 2.**
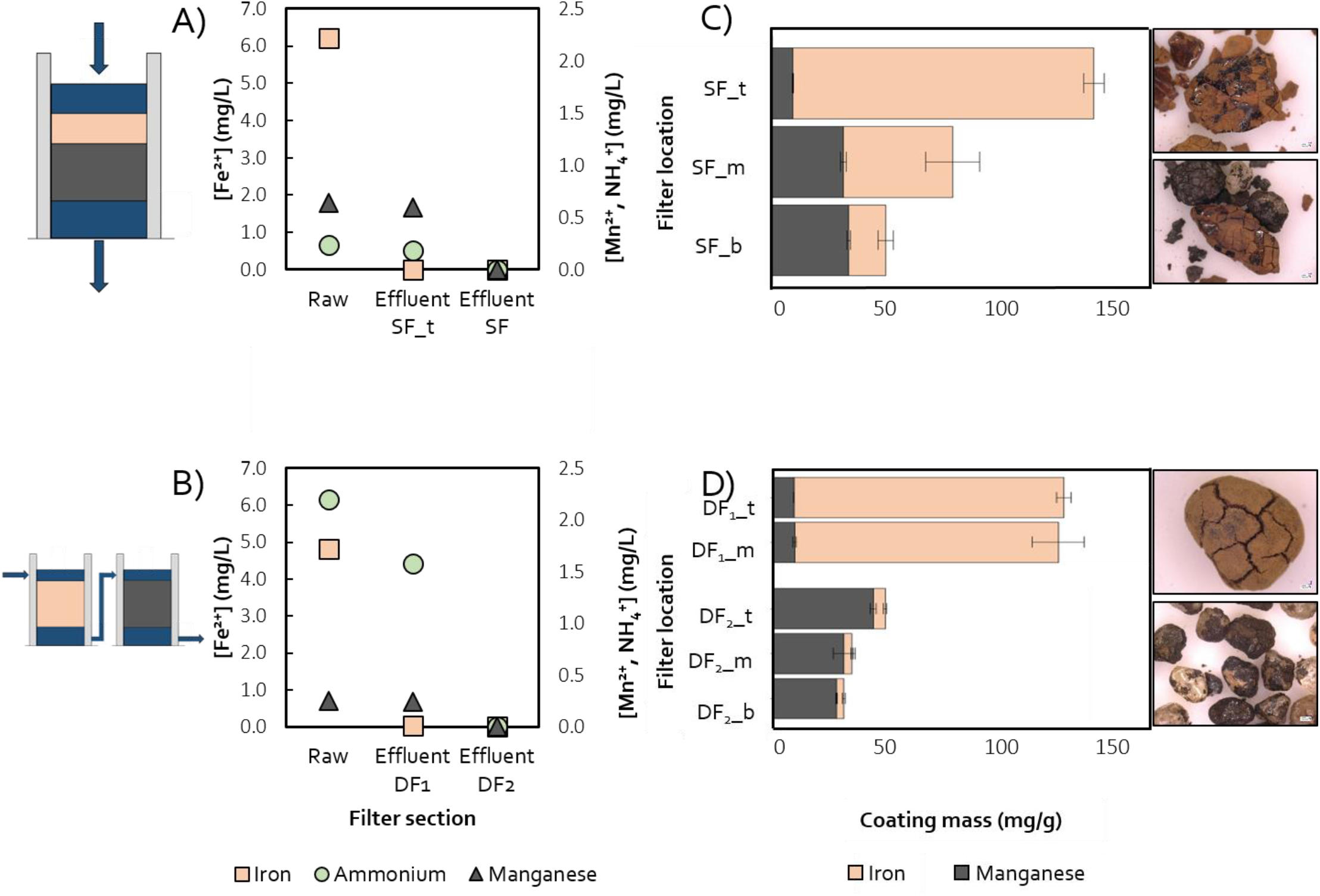
Concentration profiles of iron, manganese, and ammonium along the rapid sand filter heights in both the (A) single- and (B) double-filter DWTPs. Total removal is achieved at the end of the treatment. Coating composition of filter medium from the (C) single- and (D) double-filter DWTPs, with microscope images of SF_t, SF_b, DF1_t and DF2_b. Measurements were carried out in duplicates (A, B) or triplicates (C, D); error bars represent standard deviation.

Overall, concentration profiles and coating composition results revealed similar compartmentalization of removal processes in the two DWTPs, irrespective of dual-media single filter or single-media double filter configurations.

### 3.2. Ex situ *NH*_*4*_^*+*^*and Mn*^*2+*^ *removal mechanisms and rates reflect* in situ *compartmentalization*

#### 3.2.1. Manganese is chemically oxidized or adsorbed depending on filter media coating

The manganese removal capacity of the filter media in the different sections was quantified in batch activity tests (Figure 3A). Complete Mn^2+^ removal was observed with manganese-coated media from the later sections of both treatments (*i.e*., SF_m, SF_b and DF2). Contrastingly, it remained incomplete with iron-coated media from the first treatment sections (*i.e*., SF_t and DF1). To further characterize the removal mechanism, desorption experiments were performed by submerging drained media in demineralized water. Mn^2+^ desorption was observed exclusively with iron-rich media (Figure 3B), suggesting that Mn^2+^ is adsorbed by iron oxides, while it is oxidized on manganese oxide-coated particles.

**Figure 3.**
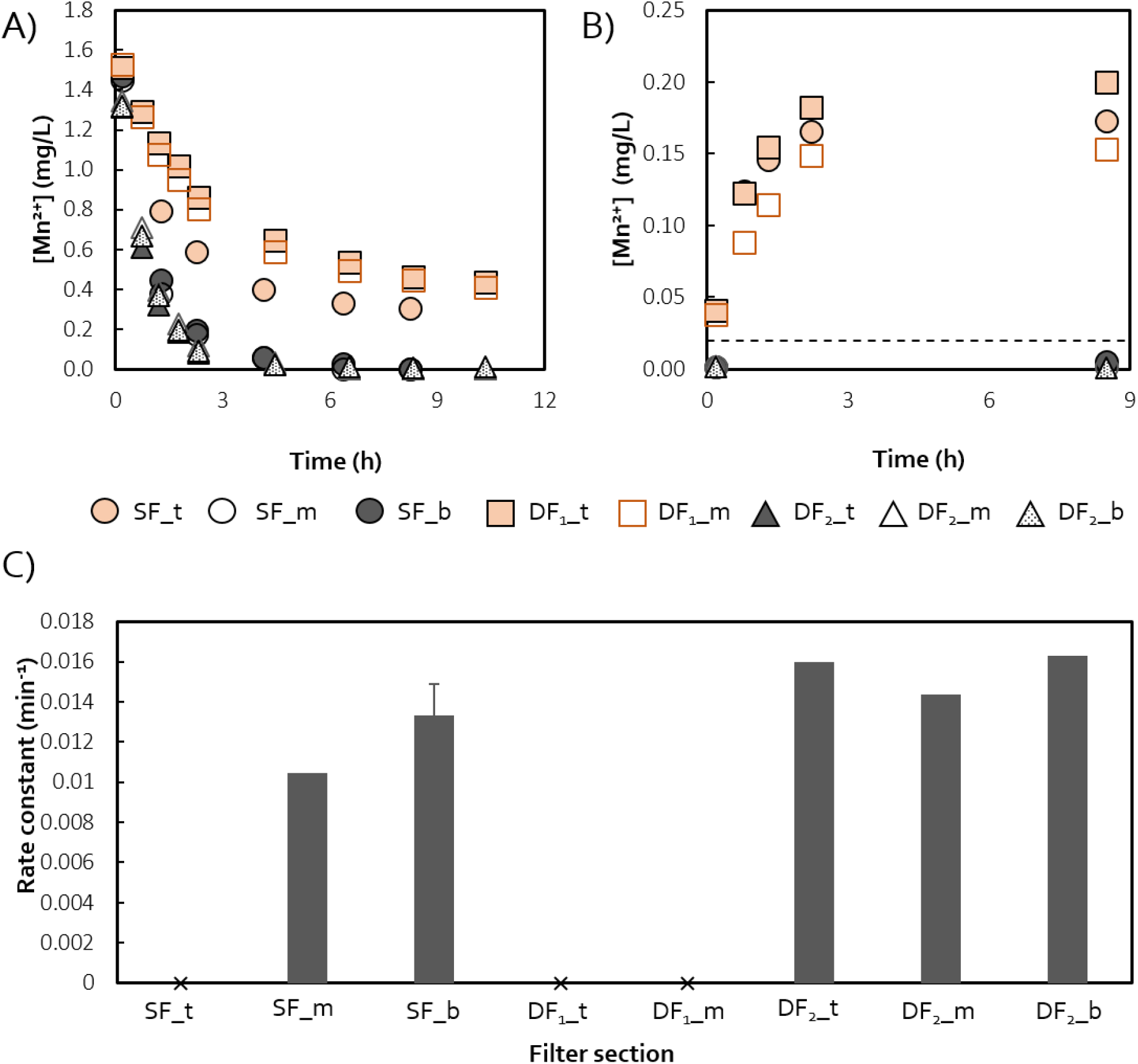
Manganese removal in (A) batch and (B) desorption tests. Colors correspond to the main component of the coating, grey for manganese and orange for iron. Batch tests were carried out in duplicate, error bars represent standard deviation (smaller than symbols). C) First-order manganese oxidation rate constants estimated with manganese-coated media. Tests were run in duplicates, values represent averages with error bars indicating standard deviation. No oxidation constants were calculated for iron-rich media where removal is primarily driven by adsorption (x).

First-order oxidation rate constants were calculated for manganese-oxidizing media (Figure 3C), as manganese oxidation is known to occur mainly physicochemically and to linearly depend on Mn^2+^ concentration (Breda, Ramsay, et al., 2019; Katsoyiannis & Zouboulis, 2004). Removal rate constants were similar in both systems (0.010-0.013 min^-1^ in SF_m and SF_b; 0.014-0.016 min^-1^ in DF2), consistent with the comparable total amounts of manganese oxide coating, the catalyst of manganese oxidation (Breda, Ramsay, et al., 2019). Moreover, the rate constants were comparable in all sections devoid of iron oxides, despite decreasing manganese concentrations along the filter height (Fig. 2). Likely, this is the result of media mixing during frequent backwash events.

#### 3.2.2. Ammonia oxidation is biological, compartmentalized and stratified within each compartment

The maximum volumetric ammonia oxidation rates along the filter heights were tested *ex situ* in batch experiments (Figure 4A). The biological nature of ammonia oxidation was validated with penicillin-amended batch tests (Figure 4B). Consistent with the measured ammonium concentration profiles along the two treatments (Figure 2A and 2B), the maximum ammonium removal rates were significantly lower in the sections where iron is present (4.5-4.7 g NH_4_^+^/(m_media_^3^·h); SF_t, DF1_t, and DF1_m) compared to the downstream sections where iron was depleted (15-62 g NH_4_^+^/(m_media_^3^·h); SF_m, SF_b, and throughout DF2). Moreover, in contrast to the relatively uniform chemical manganese oxidation rate constants (Figure 3C), the biological ammonium removal rates displayed a decreasing trend, likely reflecting the decreasing substrate availability along the filter height (Figure 4A).

**Figure 4.**
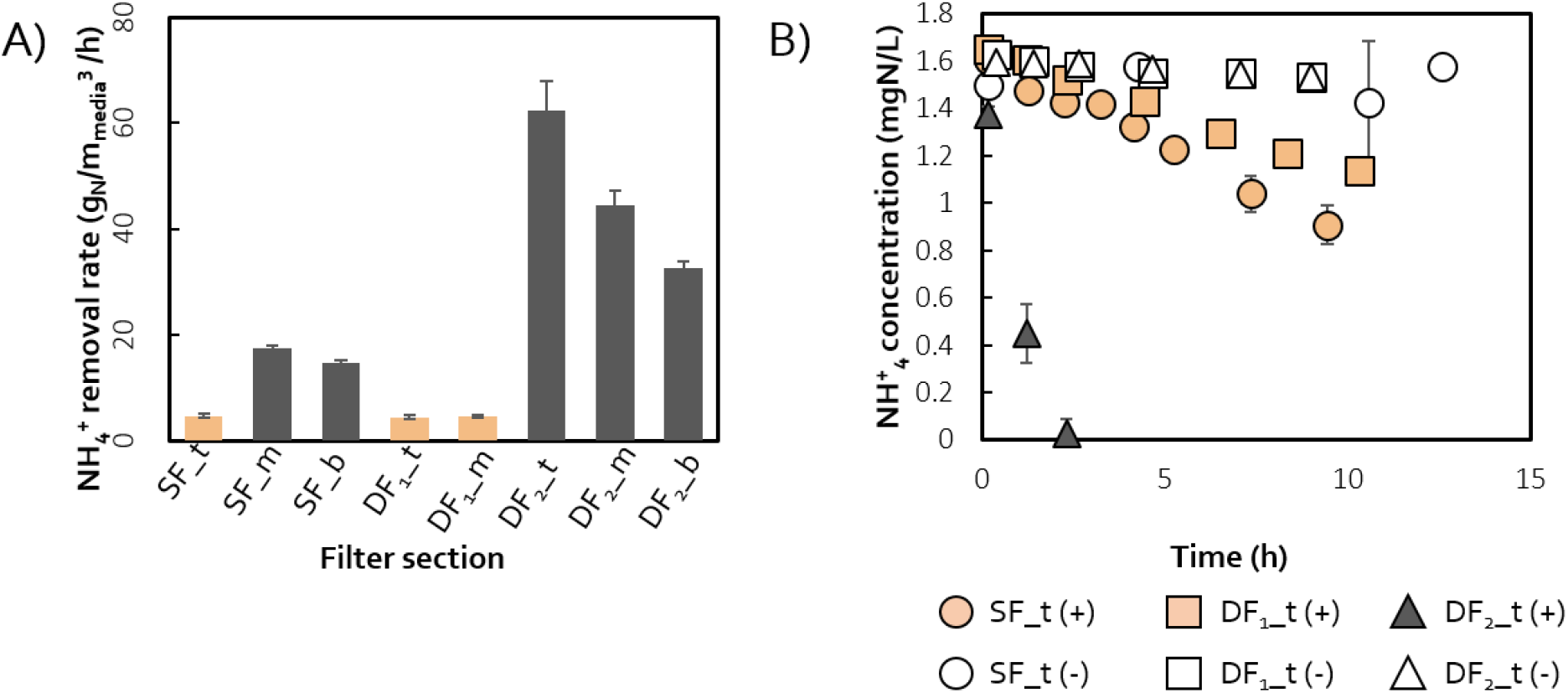
A) Maximum volumetric ammonium removal rates of the filter medium at each filter section. B) Ammonium concentration profiles in batch tests with fresh (full symbols) and penicillin-inactivated (empty symbols) filter media from the top section of each filter. All activities were quantified in triplicate, error bars represent standard deviation. The color of the symbols corresponds to the main component of the media coating, iron in orange and manganese in grey.

### 3.3. Taxonomic and functional analysis bridge plant operation with microbial physiology

#### 3.3.1. Metagenomic-based microbial composition reflects process compartmentalization

The metagenome of the microbial community in each section was sequenced to characterize the phylum-level taxonomic profile of each RSF, and to serve as reference database for metaproteomics. After quality filtering, an average of 7·10^6^ ± 3.1·10^6^ paired-end reads were obtained per sample (Table S1). *De novo* assembly of trimmed reads yielded 1·10^5^ ± 5.8·10^4^ contigs (≥ 0.5 kb) with an average N50 of 1118 ± 215 bases. Samples from the lower sections of each treatment, namely SF_m, SF_b and DF2_b, yielded the shortest contigs with an average length < 1 kb. An average of 1.8·10^6^ ± 1·10^6^ coding sequences per sample were predicted on the assembled contigs.

*Bacteria* accounted for 91.2 ± 1.9 % of the total contigs in all samples, while *Archaea* accounted for < 1% and the remainder remained unclassified. 75.4 ± 3.7 of the bacterial contigs were classified at phylum level (Figure 5). *Proteobacteria* (45.7 ± 12.1 %) and *Nitrospirae* (18.8 ± 10.8 %) dominated all sections, followed by *Planctomycetes* and *Acidobacteria* that were present in all sections except for SF_t. *Verrucomicrobia* and *Bacteroidetes* were found in SF_t and DF1, and *Actinobacteria* only in DF1. Importantly, contigs associated with the *Nitrosomonadaceae* and *Nitrospiraceae* families (Figure S2) were more abundant in the upper parts of the filters where the actual removal of ammonium occurred. In analogy, contigs assigned to phyla or families known to contribute to iron oxidation were present in all compartments but were mainly present in the anthracite layer of SF, where iron removal occurs. Overall, the composition of the microbial communities reflected the observed activity compartmentalization (Fig. 2 and 4), and this was especially evident in the dual filter system featuring a uniform community composition within each filter (*i.e*., DF1 and DF2). Despite the potential mixing effect of backwash, an analogous compartmentalization was also observed between the anthracite (SF_t) and sand layers (SF_m, SF_b) in SF.

**Figure 5.**
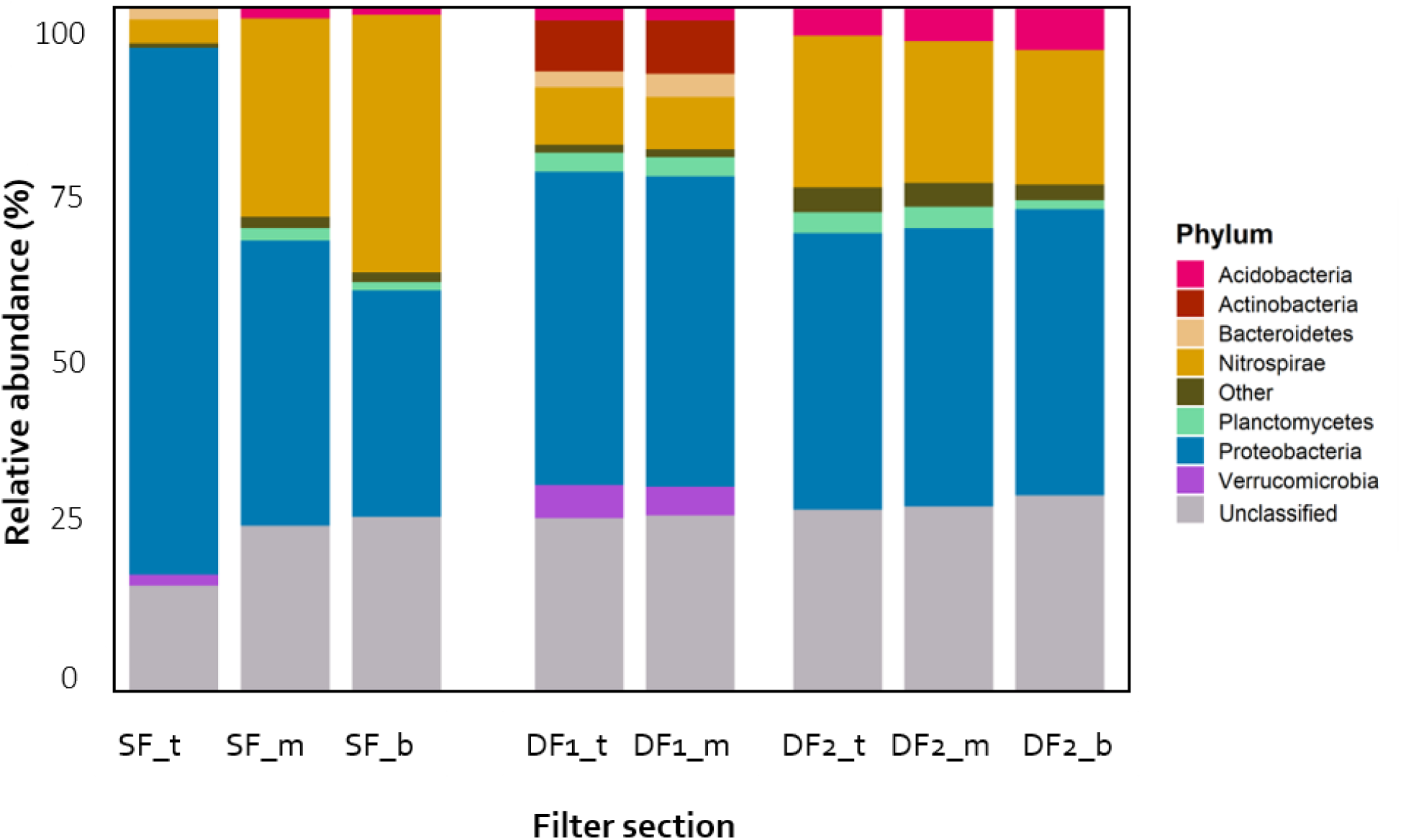
Phylum-level taxonomic classification and relative abundance calculated as percentage of total contigs of the most abundant (> 1 %) phyla in each filter section. The taxonomic classification of each contig was performed using the Contig Annotation Tool (Von Meijenfeldt et al., 2019b) with standard parameters using contigs as input and the NCBI reference database.

#### 3.3.2. Differential protein abundance explains biological activity stratification

The proteome of each section was quantified to estimate the relative contribution of known nitrogen and iron metabolizing guilds to the total biomass, as well as the distribution of core nitrogen, iron and manganese transforming enzymes. Per sample, we obtained on average 176 ± 79 individual protein groups that contained at least two unique peptides and false discover rate (FDR) < 1% (Table S4). The guilds of interest were identified by taxonomically grouping all proteins at genus level, and subsequently classifying the resulting genera based on their most-probable energy source, *i.e*., iron, nitrogen or “others” (Figure 6 and Table S3) based on the reported physiologies of the respective genus. The relative contribution of individual proteins of interest was estimated by grouping all proteins with the same KO number and summing up all the corresponding relative abundances.

**Figure 6.**
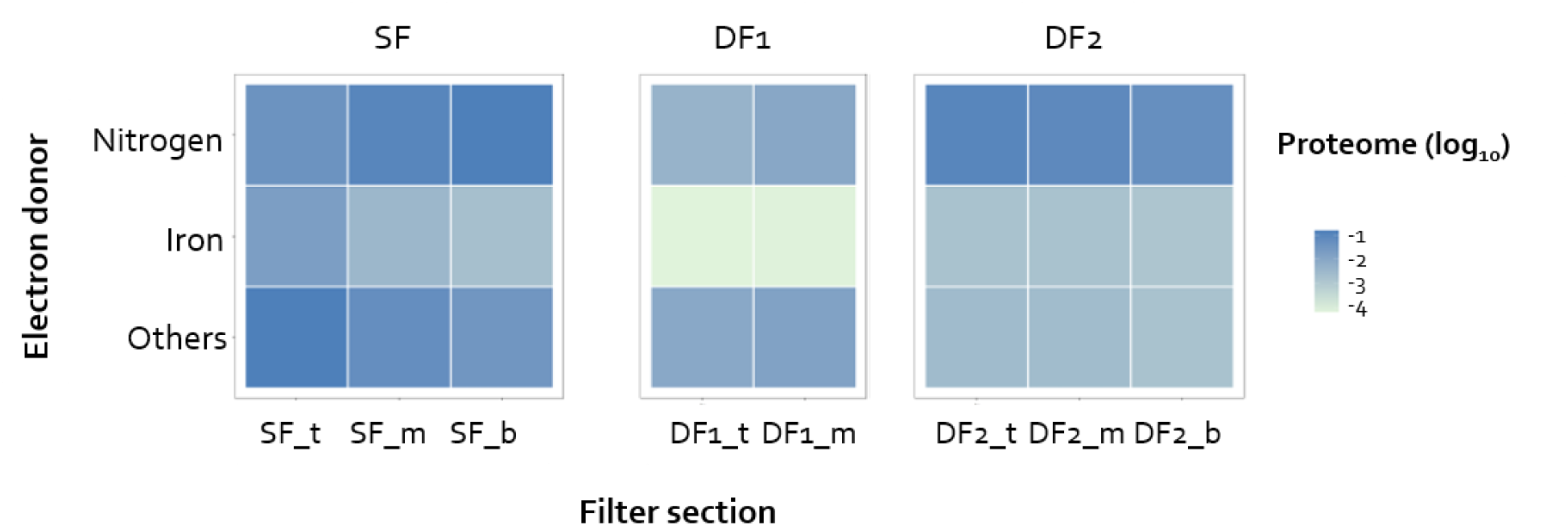
Relative protein-based biomass abundance expressed as relative spectral counts (*i.e*., protein weight normalized spectral counts divided by the sum of spectral counts of all proteins of the 8 samples together) of iron, nitrogen and ‘others’ cycling organisms, based on the most-probable energy source at genus level. Only genera with at least 2 different identified proteins (hits) were used. No genus directly associated with manganese oxidation was found. Detailed information can be found in Supplementary Table 3.

The protein-based quantification of nitrogen metabolizing guilds revealed a clear increase in their relative abundance in the lower sections of SF and in DF2, with a marked stratification in DF2 (Figure 6), in analogy with the *ex-situ* activities (Figure 4A). The genes of core nitrogen metabolisms - nitrification, denitrification, and dissimilatory nitrite reduction to ammonium (DNRA) - were present in all samples (Figure 7A; Table S2), and the abundance of the corresponding proteins was higher in the downstream sections of both systems. Specifically, the key proteins for nitrification, ammonia monooxygenase (AmoABC) and nitrite oxidoreductase (NxrAB), were present in all samples. Conversely, hydroxylamine dehydrogenase (Hao) was found at much lower abundances and only in some sections, while its coding gene *hao* was ubiquitously present. In terms of denitrification, nitrite reductase (NirK or NirS) was the only detected protein even if most genes, *i.e*., nitrate reductase (*narGHI*), nitric oxide reductase (*norBC*) and nitrous oxide reductase (*nosZ*), were found in the metagenome of all sections. In analogy, DNRA genes were found in most metagenomes but none of the corresponding proteins were detected (data not shown; Table S2).

**Figure 7.**
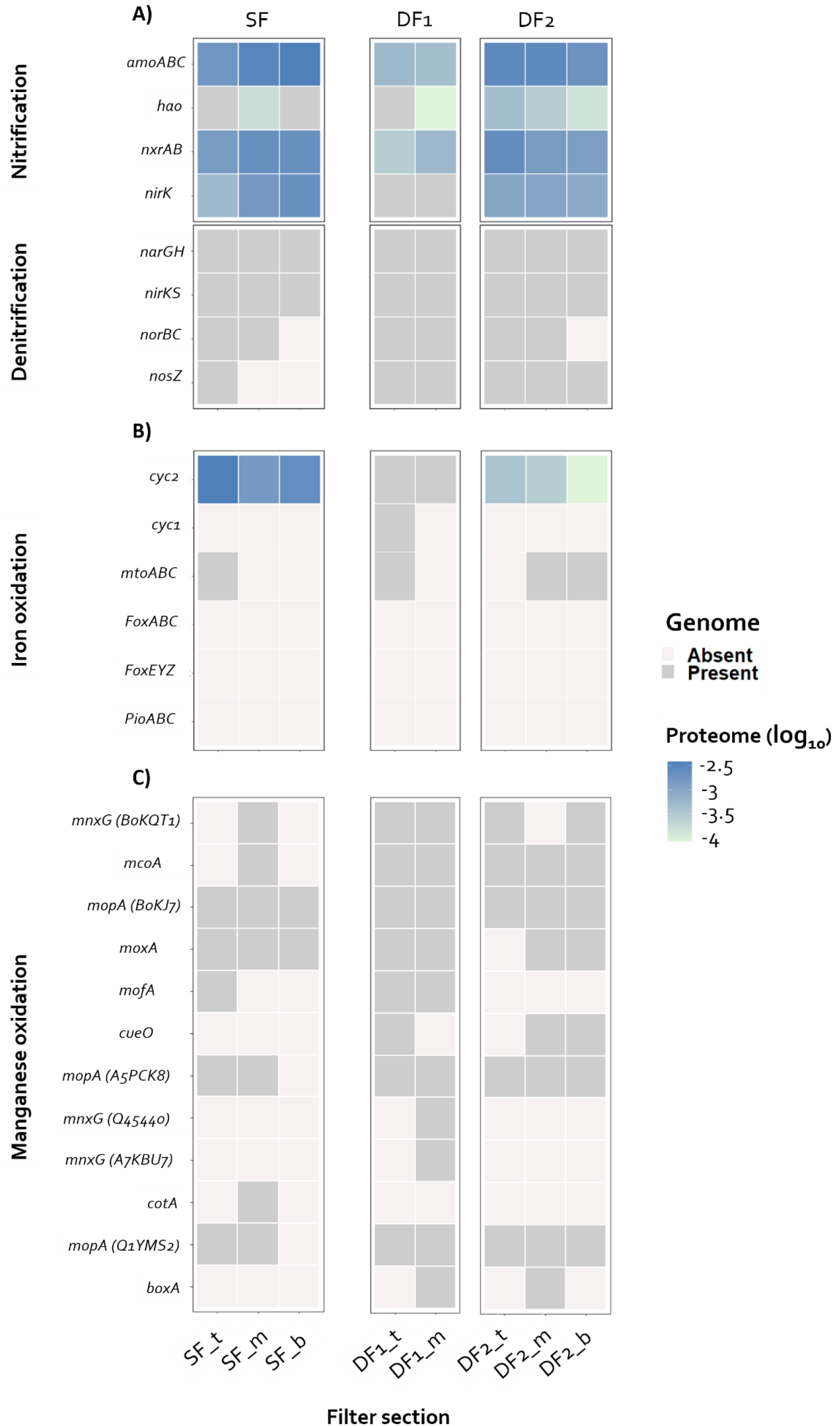
Relative protein abundance (protein-mass normalized spectral counts divided by the total spectral counts of the 8 samples) of the main (A) nitrogen, (B) iron and (C) manganese metabolism genes. Genes were selected based on the nitrification and denitrification pathways (#M00528, #M00529) of the (A) Kyoto Encyclopedia of Genes and Genomes, (B) FeGenie (Garber et al., 2020), and (C) a custom-made database based on (Hu et al., 2020). Dark grey, genes found at least once in the metagenome but not in the metaproteome; light grey, genes not found in the metagenome. Detailed information about each enzyme can be found in Supplementary Table 2.

Consistent with the iron concentration profiles and the coating composition along SF (Figure 2), putative iron-oxidizers were most abundant in the first section of SF (SF_t; Figure 6). In turn, surprisingly low abundance of iron-oxidizers was found in DF1, despite all iron being removed in this filter. This observation coincides with low overall biomass abundances in DF1_t and DF1_m, calculated as the sum of normalized spectral counts of each section divided by the sum of spectral counts of all samples combined, as compared to the other filters (Figure S3). Genes encoding for putative iron oxidizing proteins Cyc2, Cyd1, MtoABC (based on FeGenie (Garber et al., 2020)) were present in all sections, yet the corresponding proteins were only found in SF and DF2 (Figure 7B). Other putative iron-oxidizing genes, *i.e*., *foxABC, foxEYZ* and *pioABC* (Garber et al., 2020), were not detected in any sample. Lastly, putative manganese-oxidizing genes were spread across all samples with no evident correlation with filter operation, but no expressed proteins were detected.

## 4. Discussion

Two full-scale drinking water treatment plants were chosen to unravel the interaction between physicochemical and biological processes, and their distribution along the height of rapid sand filters (RSFs). While treating groundwater with similar compositions, the first plant employed a single-filter with an anthracite layer on top of a quartz sand core, and the second comprised a double-filter configuration containing quartz sand only. *In situ* and *ex situ* quantification of iron, manganese, and ammonium removals were combined with the characterization of the corresponding filter-media coating. Genomic and proteomic profiling along the filter height was used to compare the metabolic potential (*i.e*., genes encoding for certain proteins) with the actual metabolism (*i.e*., expressed proteins) of the underlying microbial communities.

### 4.1. Backwash-driven filter-media mixing results in compartments with homogeneous coating and genetic fingerprints

Both sand filtration systems exhibited highly comparable operational performances and harboured microbial communities with surprisingly similar taxonomic and functional profiles. Iron was removed in the first sections, being it the anthracite top layer in the single filter system or the first filter in the double filter configuration, while ammonium and manganese removal occurred primarily in the downstream sections (Figure 2).

Spatial separation between iron and manganese removal has often been reported in rapid sand filters (Bourgine et al., 1994; Gouzinis et al., 1998), and it is commonly attributed to the different pH/oxidation-reduction potentials required for their oxidation and precipitation (Pacini et al., 2005). Also, the mineral coating composition is often used as a proxy to localize where removals occur during the lifespan of a filter (Gülay et al., 2014). Consistently, we observed the process compartmentalization clearly reflected in the coating composition of the filter media of both studied systems. Iron and manganese oxides dominated the first and the downstream sections of each system, respectively (Figure 2), with coating amounts comparable to other DWTPs (Breda et al., 2019). Moreover, both the manganese coating mass and the removal rate constants were rather constant in all sections devoid of iron (Figure 3). As the physicochemical oxidation of manganese is directly proportional to the available concentration of manganese (Breda, Ramsay, et al., 2019; Katsoyiannis & Zouboulis, 2004), manganese oxide coating was expected to decrease along the filter height. In contrast, we observed homogeneity of coating thickness within the manganese-removing compartments of both filter systems, which is likely the result of filter-media mixing during backwash (Ramsay et al., 2021).

The observed compartmentalization of *in situ* removal rates and metal oxide coating types was also supported by the taxonomic classification of the shotgun metagenomics-derived contigs (Figure 5), and by the distribution of genes encoding for proteins involved in the oxidation of iron, ammonium, and manganese (Figure 7). The relative abundance of families commonly associated to the oxidation of iron (*Gallionellaceae*; (Gülay et al., 2018)) and nitrogen (*Nitrosomonadaceae* and *Nitrospiraceae*; (Fowler et al., 2018)) in RSFs were higher in the top and downstream sections of each DWTP, respectively (Figure S1). Moreover, their distribution within each compartment was relatively homogeneous (Figure 5) suggesting that microbial growth on the available substrates in each section is slower compared to the frequency of backwash filter media mixing. Similar spatial separation between iron and ammonium removal (Bourgine et al., 1994; de Vet et al., 2009; Sharma et al., 2005) but low heterogeneity in genome-based microbial community compositions have been observed along the height of different filters previously (Bai et al., 2013; Tatari et al., 2017). de Vet et al., (2009) ascribed this phenomenon to the accumulation of iron hydroxide flocks in the filter media, and phosphate (de Vet et al., 2012) or copper (Wagner et al., 2016) limitation have also been shown to hinder nitrification in RSFs. While the direct impact of iron oxides needs further investigation, our work allows to rule out the impact of nutrient limitations in the investigated DWTPs as full nitrification was not suppressed but simply spatially delayed.

Taken together, these results provide solid evidence that irrespective of treatment configuration, being it a dual-media single filter or single-media double filter setup, processes compartmentalize along filter heights. The fact that anthracite and quartz sand remained clearly separated in the single filter after over one year of operation also proves the effectiveness of dual-media filtration in spatially separating processes, and its equivalence to two filters in series. Moreover, the homogeneity of media coating and genome-based microbial community composition within each compartment highlights the process-wise relevance of backwashing filter media mixing.

### 4.2. Protein abundance profiles reflect biological activity stratification and substrate gradients

In stark contrast to the clear intra-compartment homogeneity, *in situ* iron, ammonium and manganese concentration profiles (Figure 2) and *ex situ* biological ammonium oxidation rates displayed a strongly stratified profile within the different filter compartments (Figure 4). To investigate the discrepancy between the stratified activities and the homogeneous metagenome-based distribution of iron-, ammonia-, and manganese-oxidizing genes (Figure 7), we employed shotgun metaproteomics to quantify the relative abundance of the encoded proteins along the filter heights as proxy for *in situ* biological activity.

Stratification patterns in *ex situ* and *in situ* ammonia oxidation rates, similar to the ones observed in SF and DF, have been reported in sand filters treating water with (Tatari et al., 2013, 2016) and without (Lee et al., 2014) iron, and have been attributed to the progressive decrease in ammonium concentration along the filter height (de Vet, Rietveld, et al., 2009). We detected higher protein-based relative abundances of putative nitrifying microorganisms in the downstream sections where most ammonia oxidation occurs, and we even observed a clear stratification in DF2 where all three investigated filter sections were devoid of iron (Figure 6). Importantly, the same decreasing trend could also be observed in the abundance of the core enzymes of nitrification, *i.e*., ammonia monooxygenase (AmoABC), nitrite oxidoreductase (NxrAB) and nitrifier-assigned nitrite reductase (NirK), used here as proxy for the actual nitrifying activity of the microbial community (Figure 7). In analogy, the protein-based relative abundance of putative iron-oxidizing bacteria was the highest in the anthracite layer of SF (Figure 6), the only section where contigs of the *Gallionellaceae* family, the most common iron-oxidizing bacteria (de Vet et al., 2011), were found. In terms of specific enzymes, the only detected Fe-oxidizing protein with a proven function was the *c*-type cytochrome Cyc2 (Castelle et al., 2008), a protein reported to be conserved in most neutrophilic iron oxidizers (McAllister et al., 2020). No other iron-oxidizing protein was detected and, likely due to the still limited biochemical characterization of this process, only few of the known iron-oxidizing genes reviewed by Garber et al. (2020) were found. The surprisingly low relative abundance of proteins assigned to putative iron oxidizers in DF1 (Figure 6) could in principle suggest a dominance of abiotic processes controlling iron oxidation in this filter. However, the overall protein recovery in DF1 was significantly lower than in all other sections. This could possibly be ascribed to iron oxides hindering protein extraction, as seen by Barco & Edwards (2014), yet this was not encountered in the iron-removing SF_t section. Thus, the reason for the low protein yield in DF1 as well as the extend of contribution of biological iron oxidation to overall iron removal deserve further investigation. Ultimately, the absence of proteins involved in manganese oxidation, despite the presence of the corresponding genes, suggests an abiotic-dominated process in strong agreement with the homogeneously distributed, first-order rate constants (Figure 3).

In conclusion, the quantification of expressed proteins allowed us to resolve the often reported dichotomy between, on the one hand, the intra-compartment homogeneity of the mineral coating and genome-based microbial community composition and, on the other, the marked stratification of the concentration profiles and biological activities. We hypothesize that backwashing results in mixing and homogenization of the filter media and thereby exposes the attached microorganisms to a different substrate load. In response, microorganisms adjust their metabolic fluxes by adapting their protein pool at a faster rate than the mixing frequency. Conversely, the actual microbial growth between mixing is minimal and does not result in detectable changes at the genome level. Beyond rapid sand filters, these results prove the potential of metagenome-guided metaproteomics as a powerful tool to understand metabolic interactions in highly dynamic ecosystems.

## 5. Conclusions

In this work, we compared two full-scale drinking water treatment plants to resolve the interaction between the physicochemical and biological processes controlling ammonium, iron and manganese removal. The first plant employed a dual-bed (anthracite and quartz sand) single filter, the second two single-bed quartz sand filters in series. Both plants treated groundwater with similar chemical composition. The combination of biological activity measurements, mineral coating characterization, and multi-omics profiling led to the following conclusions:

^*^both systems exhibited comparable performances and clear process compartmentalization, with iron being fully removed in the first, and ammonium and manganese primarily in the downstream sections;

^*^the mineral composition of the media coating reflected the compartmentalization of processes, *i.e*., iron and manganese oxides dominated the first and downstream sections of each system, respectively;

^*^the homogeneity of media coating and genome-based composition of the microbial communities within each compartment highlighted the process-wise relevance of backwashing filter media mixing;

^*^only protein abundance profiles reflected the intra-compartment removal stratification, strongly highlighting the potential of metaproteomic approaches for studying highly dynamic ecosystems;

^*^dual-media single filters and series of single-media filters are equivalent from an operational and ecological perspective, and anthracite and sand remain separated despite frequent backwashing.

## List of abbreviations

RSF: rapid sand filter
DWTP: drinking water treatment plant
SF: single filter
DF: double filter
DNRA: dissimilatory nitrate reduction to ammonia

## 6. Contributions

FCR, ML, and MvL and DvH conceived the study. FCR, ML, NK and MvL and DvH designed the experiments. FCR and NK performed the experiments with contributions from FS (field work), SM (mineral coating) and TvA (metagenomic sequencing). MP conducted the metaproteomic analysis. FCR, ML, NK, SL, MP, MvL and DvH performed data analysis and/or helped interpret the results. FCR wrote the manuscript, with contributions from ML, NK, SL, MP, MvL and DvH. All co-authors critically reviewed the manuscript and approved the final version.

## 7. Acknowledgements

The authors would like to thank dr. Jaap Dijkshoorn (Vitens N.V.) for their fieldwork help and insightful discussions, ir. Carol de Ram for the metaproteomic analysis and ir. Nina Roothans for establishing a pipeline for metagenomics.

This work was financed by the NWO partnership program ‘Dunea–Vitens: Sand Filtration’ (projects 17830 and 17841) of the Dutch Research Council (NWO) and the drinking water companies Vitens NV and Dunea Duin & Water. ML and SL were supported by NWO (project numbers VI.Veni.192.252 and 016.Vidi.189.050, respectively).

