## Supplementary Material for "Meta-omics profiling of full-scale groundwater rapid sand filters explains stratification of iron, ammonium and manganese removals"

### 1. Supplementary information

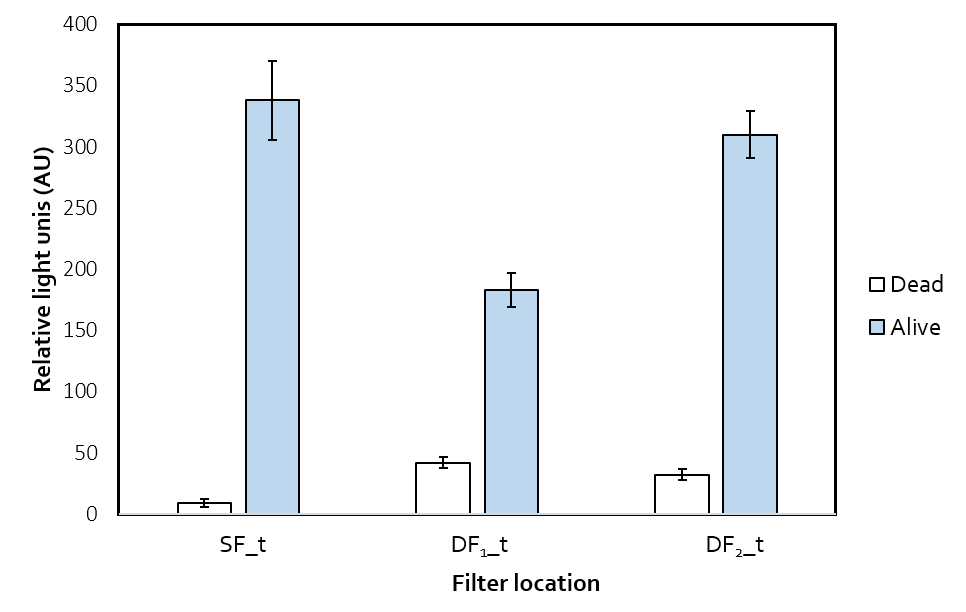

Figure S1. ATP concentration from filter media used in ammonium conversion activity tests as relative light units (AU). Blue bars correspond to fresh media, white bars to penicillin-inactivated media. Vertical bars represent standard deviations from experimental triplicate measurements.

Table S1. Sequencing and assembly statistics for all samples.

| **Statistics** | **Single-filter (SF)** | | | **Double-filter 1 (DF1)** | | **Double-filter 2 (DF2)** | | |
| --- | --- | --- | --- | --- | --- | --- | --- | --- |
|  | *SF_t* | *SF_m* | *SF_b* | *DF1_t* | *DF1_m* | *DF2_t* | *DF2_m* | *DF2_b* |
| Raw reads | 6.24E+06 | 2.93E+06 | 1.37E+06 | 8.42E+06 | 8.27E+06 | 1.04E+07 | 8.83E+06 | 9.68E+06 |
| Trimmed reads | 6.22E+06 | 2.82E+06 | 1.32E+06 | 8.40E+06 | 8.24E+06 | 8.40E+06 | 8.77E+06 | 4.93E+06 |
| Contigs | 4.27E+04 | 3.68E+04 | 1.38E+04 | 1.45E+05 | 1.46E+05 | 1.68E+05 | 1.61E+05 | 1.01E+05 |
| Average contig size | 1273 | 967 | 841 | 1187 | 1200 | 1075 | 1041 | 990 |
| N50 | 1444 | 953 | 783 | 1323 | 1353 | 1074 | 1036 | 981 |
| Total bp in contigs | 5.44E+07 | 3.56E+07 | 1.16E+07 | 1.73E+08 | 1.75E+08 | 1.80E+08 | 1.67E+08 | 1.00E+08 |
| Longest contig | 1.07E+05 | 9.13E+04 | 1.09E+05 | 1.32E+05 | 1.44E+05 | 1.57E+05 | 9.20E+04 | 6.76E+04 |
| % reads mapped to contigs | 88.8 | 54.6 | 43.7 | 83.6 | 83.8 | 79.9 | 77.6 | 67.7 |

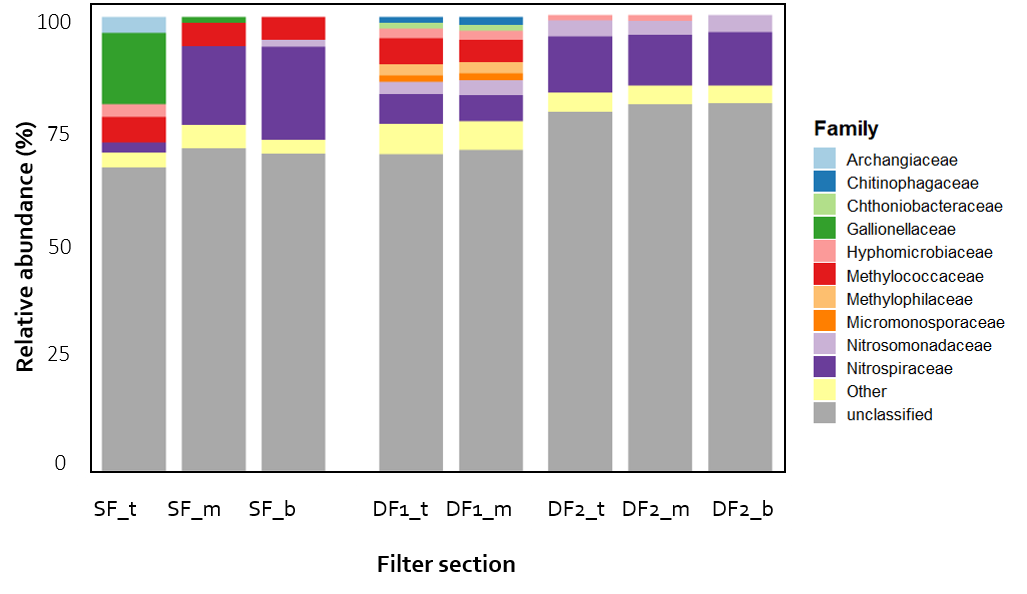

Figure S2. Family-level taxonomic classification and relative abundance of the most abundant (> 1 %) families in each filter section. The taxonomic classification of each contig was performed using the Contig Annotation Tool (Von Meijenfeldt et al., 2019b) with standard parameters using the NCBI reference database.

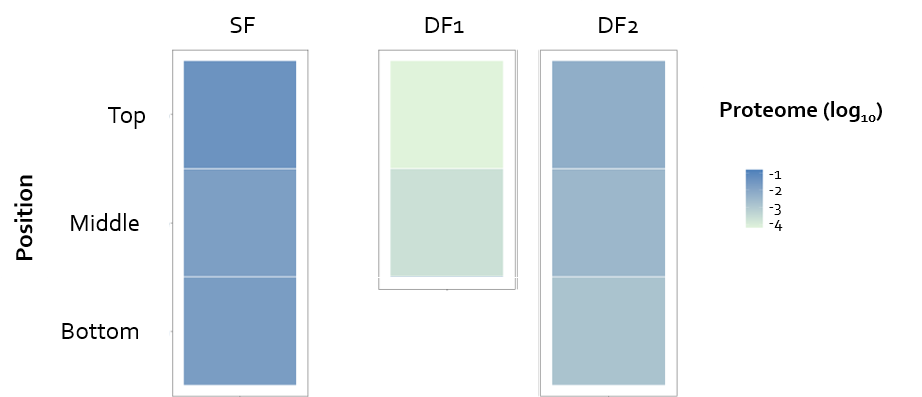

Figure S3. Relative protein-based biomass abundance as normalized spectral counts per filter section.

Table S2. Main nitrification, denitrification, DNRA, iron oxidation and manganese oxidation genes present (gray) or absent (white) in the metagenome of each sample. Gene annotation was carried out with GhostKOALA for nitrogen metabolism, FeGenie (Garber et al., 2020) for iron oxidation and blastp employing a custom-made database for manganese oxidation.

(See attached Excel sheet)

Table S3. Classification of the 20 most abundant genera identified with metaproteomics based on their most probable energy metabolism.

| **Genus** | **Energy source** | **Reference** |
| --- | --- | --- |
| *Candidatus* Brocadia | Nitrogen | Schmid et al, 2001 |
| *Candidatus* Methylospira | Others | Danilova et al, 2016 |
| *Candidatus* Nitrotoga | Nitrogen | Kitzinger et al, 2018 |
| Caulobacteraceae bacterium OTSz_A_272 | Others | Braun and Szewzyk, 2016 |
| *Ferrigenium* | Iron | Khalifa et al, 2018 |
| *Filomicrobium* | Iron | Schlesner, 1987 |
| *Gallionella* | Iron | de Vet, 2011 |
| *Hyphomicrobium* | Others | Rainey, 1998 |
| *Methylobacillus* | Others | Yordy and Weaver, 1977 |
| *Methyloglobus* | Others | Deutzmann et al, 2014 |
| *Methylomonas* | Others | Bowman et al, 1993 |
| *Methylotuvimicrobium* | Others | Orata et al, 2018 |
| *Methylovorus* | Others | Doronina et al, 2005 |
| *Methylovulum* | Others | Oshkin et al, 2016 |
| *Nitrosomonas* | Nitrogen | Winogradsky, 1892 |
| *Nitrosospira* | Nitrogen | Winogradsky S & Winogradsky H, 1933 |
| *Nitrospira* | Nitrogen | Watson et al, 1986 |
| *Sideroxydans* | Iron | Emerson et al, 2013 |

Table S4. Filter media density at each location. Density was determined in triplicate using the Archimedes principle: wet sand was added to a measuring cylinder containing water while tracking mass and volume increase. The measurements were carried out in triplicate, with error represented as standard deviation.

| Location | Density (g·cm^-3^) |
| --- | --- |
| SF_t | 1.28 ± 0.02 |
| SF_m | 1.69 ± 0.02 |
| SF_B | 1.72 ± 0.01 |
| DF1_t | 1.73 ± 0.04 |
| DF1_m | 1.67 ± 0.03 |
| DF2_t | 1.68 ± 0.04 |
| DF2_m | 1.73 ± 0.04 |
| DF2_b | 1.77 ± 0.03 |

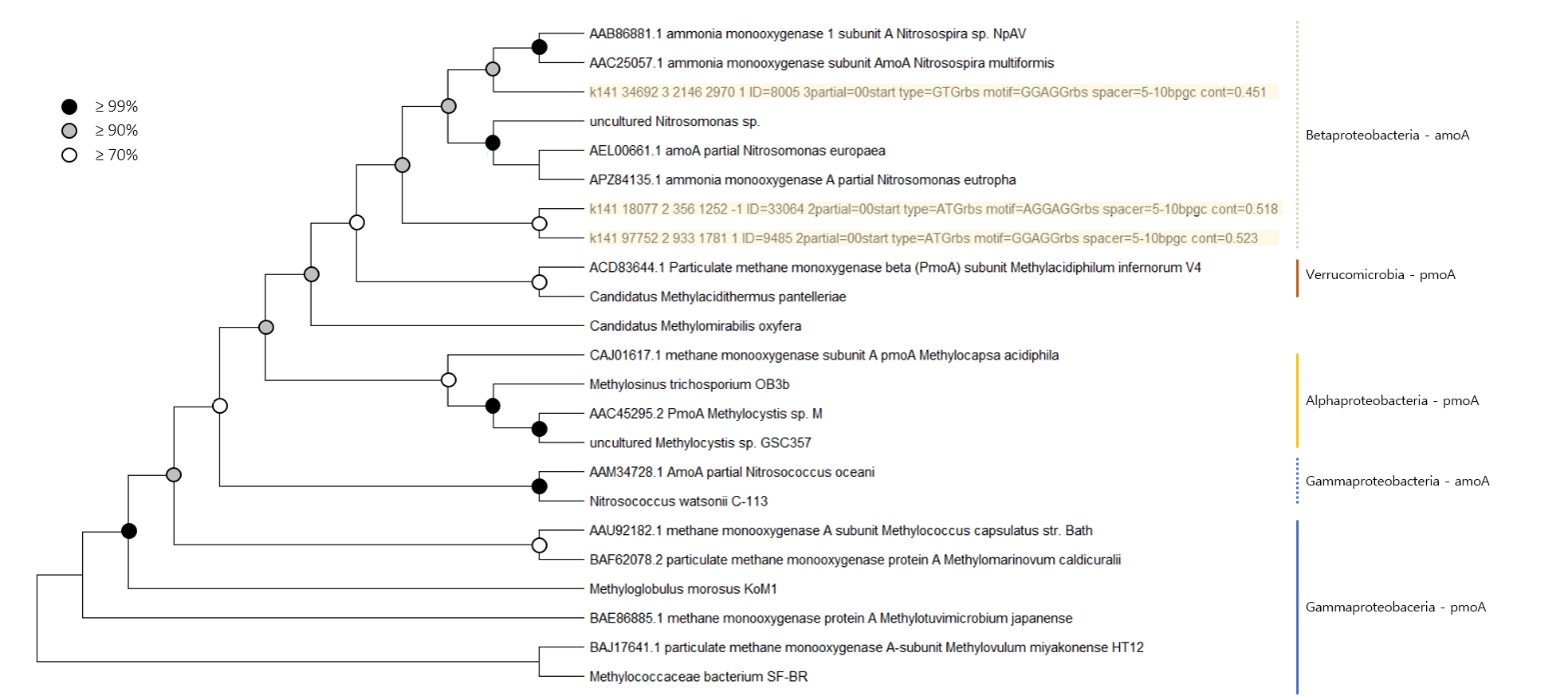

Figure S4. Bootstrapped phylogenetic tree with protein sequences of ammonia and methane monooxygenase subunit A. The tree indicated the affiliation of the three proteins identified in this study, with their contig number as name. Reference sequences were obtained from NCBI. White, grey, and black dots represent bootstrap support ≥ 70, 90 and 99 %, respectively.

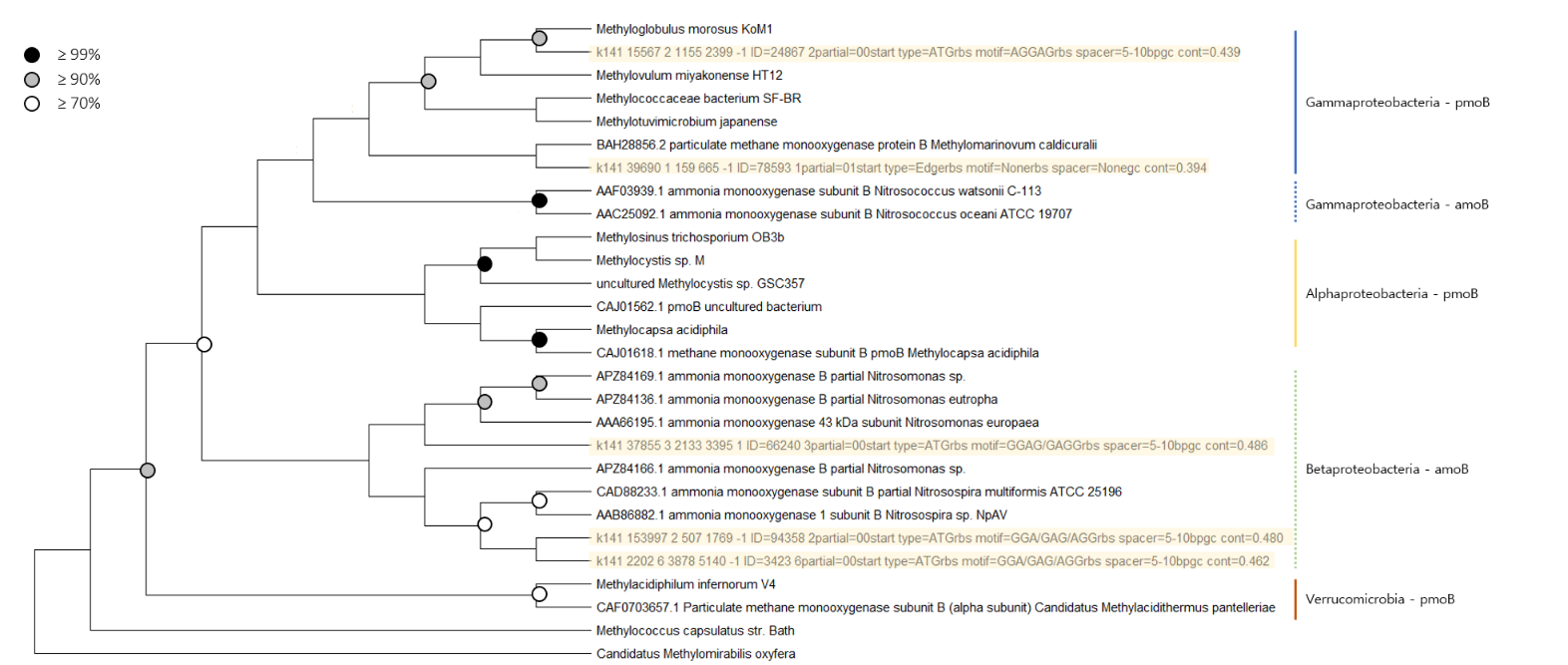

Figure S5. Bootstrapped phylogenetic tree with protein sequences of ammonia and methane monooxygenase subunit B. The tree indicates the affiliation of the five identified proteins, with their contig number as name. Reference sequences were obtained from NCBI. White, grey, and black dots represent bootstraps ≥ 70, 90 and 99 %, respectively.

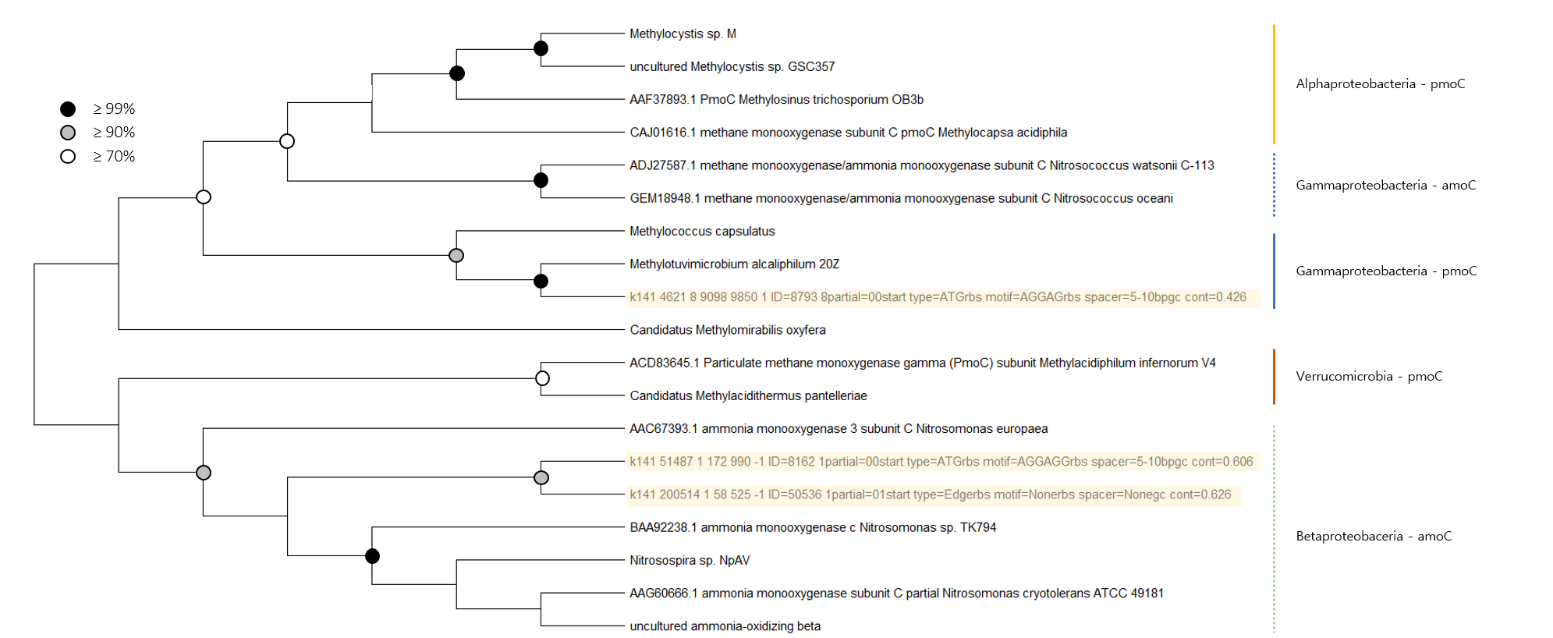

Figure S6. Bootstrapped phylogenetic tree with protein sequences of ammonia and methane monooxygenase subunit C. The tree indicates the affiliation of the five identified proteins, with their contig number as name. Reference sequences were obtained from NCBI. White, grey, and black dots represent bootstraps ≥ 70, 90 and 99 %, respectively.

Table S5. Raw metaproteomic data

See attached Excel Sheet

##### **References supplementary material**

Albertsen, M., Hugenholtz, P., Skarshewski, A., Nielsen, K. L., Tyson, G. W., & Nielsen, P. H. (2013). Genome sequences of rare, uncultured bacteria obtained by differential coverage binning of multiple metagenomes. *Nature Biotechnology*, *31*(6), 533–538. https://doi.org/10.1038/nbt.2579

Andrews, S. (2010). *FastQC: A Quality Control Tool for High Throughput Sequence Data [Online]*. http://www.bioinformatics.babraham.ac.uk/projects/fastqc/

APHA. (2018). *3500-Fe IRON - Standard Methods For the Examination of Water and Wastewater*. https://www.standardmethods.org/doi/full/10.2105/SMWW.2882.055

Bai, Y., Liu, R., Liang, J., & Qu, J. (2013). Integrated Metagenomic and Physiochemical Analyses to Evaluate the Potential Role of Microbes in the Sand Filter of a Drinking Water Treatment System. *PLoS ONE*, *8*(4). https://doi.org/10.1371/journal.pone.0061011

Barco, R. A., & Edwards, K. J. (2014). Interactions of proteins with biogenic iron oxyhydroxides and a new culturing technique to increase biomass yields of neutrophilic, iron-oxidizing bacteria. *Frontiers in Microbiology*, *5*(MAY), 259. https://doi.org/10.3389/FMICB.2014.00259/ABSTRACT

Bolger, A. M., Lohse, M., & Usadel, B. (2014). Trimmomatic: a flexible trimmer for Illumina sequence data. *Bioinformatics (Oxford, England)*, *30*(15), 2114–2120. https://doi.org/10.1093/BIOINFORMATICS/BTU170

Bourgine, F. P., Gennery, M., Chapman, J. I., Kerai, H., Green, J. G., Rap, R. J., Ellis, S., & Gaumard, C. (1994). Biological Processes at Saints Hill Water‐Treatment Plant, Kent. *Water and Environment Journal*, *8*(4), 379–391. https://doi.org/10.1111/j.1747-6593.1994.tb01121.x

Breda, I. L., Ramsay, L., Søborg, D. A., Dimitrova, R., & Roslev, P. (2019). Manganese removal processes at 10 groundwater fed full-scale drinking water treatment plants. *Water Quality Research Journal of Canada*, *54*(4), 326–337. https://doi.org/10.2166/wqrj.2019.006

Breda, I. L., Søborg, D. A., Ramsay, L., & Roslev, P. (2019). Manganese removal processes during start-up of inoculated and non-inoculated drinking water biofilters. *Water Quality Research Journal of Canada*, *54*(1), 47–56. https://doi.org/10.2166/wqrj.2018.016

Bruins, J. H., Petrusevski, B., Slokar, Y. M., Wübbels, G. H., Huysman, K., Wullings, B. A., Joris, K., Kruithof, J. C., & Kennedy, M. D. (2017). Identification of the bacterial population in manganese removal filters. *Water Science and Technology: Water Supply*, *17*(3), 842–850. https://doi.org/10.2166/ws.2016.184

Buchfink, B., Xie, C., & Huson, D. H. (2014). Fast and sensitive protein alignment using DIAMOND. *Nature Methods 2014 12:1*, *12*(1), 59–60. https://doi.org/10.1038/NMETH.3176

Burger, M. S., Krentz, C. A., Mercer, S. S., & Gagnon, G. A. (2008). Manganese removal and occurrence of manganese oxidizing bacteria in full-scale biofilters. *Journal of Water Supply: Research and Technology - AQUA*, *57*(5), 351–359. https://doi.org/10.2166/aqua.2008.050

Castelle, C., Guiral, M., Malarte, G., Ledgham, F., Leroy, G., Brugna, M., & Giudici-Orticoni, M. T. (2008). A new iron-oxidizing/O2-reducing supercomplex spanning both inner and outer membranes, isolated from the extreme acidophile Acidithiobacillus ferrooxidans. *Journal of Biological Chemistry*, *283*(38), 25803–25811. https://doi.org/10.1074/jbc.M802496200

Claff, S. R., Sullivan, L. A., Burton, E. D., & Bush, R. T. (2010). A sequential extraction procedure for acid sulfate soils: Partitioning of iron. *Geoderma*, *155*(3–4), 224–230. https://doi.org/10.1016/J.GEODERMA.2009.12.002

de Vet, W., Dinkla, I. J. T., Muyzer, G., Rietveld, L. C., & van Loosdrecht, M. C. M. (2009). Molecular characterization of microbial populations in groundwater sources and sand filters for drinking water production. *Water Research*, *43*(1), 182–194. https://doi.org/10.1016/j.watres.2008.09.038

de Vet, W., Dinkla, I. J. T., Rietveld, L. C., & van Loosdrecht, M. C. M. (2011). Biological iron oxidation by Gallionella spp. in drinking water production under fully aerated conditions. *Water Research*, *45*(17), 5389–5398. https://doi.org/10.1016/j.watres.2011.07.028

de Vet, W., Rietveld, L. C., & Van Loosdrecht, M. C. M. (2009). Influence of iron on nitrification in full-scale drinking water trickling filters. *Journal of Water Supply: Research and Technology - AQUA*, *58.4*, 247–256. https://doi.org/10.2166/aqua.2009.115

de Vet, W., Van Loosdrecht, M. C. M., & Rietveld, L. C. (2012). Phosphorus limitation in nitrifying groundwater filters. *Water Research*, *46*(4), 1061–1069. https://doi.org/10.1016/j.watres.2011.11.075

Fowler, S. J., Palomo, A., Dechesne, A., Mines, P. D., & Smets, B. F. (2018). Comammox Nitrospira are abundant ammonia oxidizers in diverse groundwater-fed rapid sand filter communities. *Environmental Microbiology*, *20*(3), 1002–1015. https://doi.org/10.1111/1462-2920.14033

Garber, A. I., Nealson, K. H., Okamoto, A., McAllister, S. M., Chan, C. S., Barco, R. A., & Merino, N. (2020). FeGenie: A Comprehensive Tool for the Identification of Iron Genes and Iron Gene Neighborhoods in Genome and Metagenome Assemblies. *Frontiers in Microbiology*, *11*, 37. https://doi.org/10.3389/fmicb.2020.00037

Giordano, M. (2009). Global Groundwater? Issues and Solutions. *Http://Dx.Doi.Org/10.1146/Annurev.Environ.030308.100251*, *34*, 153–178. https://doi.org/10.1146/ANNUREV.ENVIRON.030308.100251

Gouzinis, A., Kosmidis, N., Vayenas, D. V., & Lyberatos, G. (1998). Removal of Mn and simultaneous removal of NH3, Fe and Mn from potable water using a trickling filter. *Water Research*, *32*(8), 2442–2450. https://doi.org/10.1016/S0043-1354(97)00471-5

Gude, J. C. J., Rietveld, L. C., & van Halem, D. (2016). Fate of low arsenic concentrations during full-scale aeration and rapid filtration. *Water Research*, *88*, 566–574. https://doi.org/10.1016/J.WATRES.2015.10.034

Gülay, A., Çekiç, Y., Musovic, S., Albrechtsen, H. J., & Smets, B. F. (2018). Diversity of Iron Oxidizers in Groundwater-Fed Rapid Sand Filters: Evidence of Fe(II)-Dependent Growth by Curvibacter and Undibacterium spp. *Frontiers in Microbiology*, *9*, 2808. https://doi.org/10.3389/fmicb.2018.02808

Gülay, A., Musovic, S., Albrechtsen, H. J., Al-Soud, W. A., Sørensen, S. J., & Smets, B. F. (2016). Ecological patterns, diversity and core taxa of microbial communities in groundwater-fed rapid gravity filters. *ISME Journal*, *10*(9), 2209–2222. https://doi.org/10.1038/ismej.2016.16

Gülay, A., Tatari, K., Musovic, S., Mateiu, R. V., Albrechtsen, H. J., & Smets, B. F. (2014). Internal porosity of mineral coating supports microbial activity in rapid sand filters for groundwater treatment. *Applied and Environmental Microbiology*, *80*(22), 7010–7020. https://doi.org/10.1128/AEM.01959-14

Hall, B. G. (2013). Building phylogenetic trees from molecular data with MEGA. *Molecular Biology and Evolution*, *30*(5), 1229–1235. https://doi.org/10.1093/molbev/mst012

Hanson, B. T., & Madsen, E. L. (2015). In situ expression of nitrite-dependent anaerobic methane oxidation proteins by Candidatus Methylomirabilis oxyfera co-occurring with expressed anammox proteins in a contaminated aquifer. *Environmental Microbiology Reports*, *7*(2), 252–264. https://doi.org/10.1111/1758-2229.12239

Hu, W., Liang, J., Ju, F., Wang, Q., Liu, R., Bai, Y., Liu, H., & Qu, J. (2020). Metagenomics Unravels Differential Microbiome Composition and Metabolic Potential in Rapid Sand Filters Purifying Surface Water Versus Groundwater. *Environmental Science and Technology*, *54*, 5206. https://doi.org/10.1021/acs.est.9b07143

Ilbert, M., & Bonnefoy, V. (2013). Insight into the evolution of the iron oxidation pathways. *Biochimica et Biophysica Acta (BBA) - Bioenergetics*, *1827*(2), 161–175. https://doi.org/10.1016/J.BBABIO.2012.10.001

Kalyuzhnaya, M. G., Gomez, O. A., & Murrell, J. C. (2019). *The Methane-Oxidizing Bacteria ( Methanotrophs )*.

Kanehisa, M., Sato, Y., & Morishima, K. (2016). BlastKOALA and GhostKOALA: KEGG Tools for Functional Characterization of Genome and Metagenome Sequences. *Journal of Molecular Biology*, *428*(4), 726–731. https://doi.org/10.1016/J.JMB.2015.11.006

Katsanou, K., & Karapanagioti, H. K. (2019). Surface water and groundwater sources for drinking water. *Handbook of Environmental Chemistry*, *67*, 1–19. https://doi.org/10.1007/698_2017_140/FIGURES/5

Katsoyiannis, I. A., & Zouboulis, A. I. (2004). Biological treatment of Mn(II) and Fe(II) containing groundwater: Kinetic considerations and product characterization. *Water Research*, *38*(7), 1922–1932. https://doi.org/10.1016/j.watres.2004.01.014

Kihn, A., Laurent, P., & Servais, P. (2000). *Measurement of potential activity of fixed nitrifying bacteria in biological filters used in drinking water production*. 161–166.

Kleikamp, H. B. C., Grouzdev, D., Schaasberg, P., Valderen, R. van, Zwaan, R. van der, Wijgaart, R. van de, Lin, Y., Abbas, B., Pronk, M., Loosdrecht, M. C. M. van, & Pabst, M. (2022). Comparative metaproteomics demonstrates different views on the complex granular sludge microbiome. *BioRxiv*, 2022.03.07.483319. https://doi.org/10.1101/2022.03.07.483319

Kleikamp, H. B. C., Pronk, M., Tugui, C., Guedes da Silva, L., Abbas, B., Lin, Y. M., van Loosdrecht, M. C. M., & Pabst, M. (2021). Database-independent de novo metaproteomics of complex microbial communities. *Cell Systems*, *12*(5), 375-383.e5. https://doi.org/10.1016/J.CELS.2021.04.003

Kleiner, M. (2019). Metaproteomics: Much More than Measuring Gene Expression in Microbial Communities. *MSystems*, *4*(3). https://doi.org/10.1128/MSYSTEMS.00115-19/ASSET/79FEFC90-FAEC-4535-828E-FD35A282F503/ASSETS/GRAPHIC/MSYSTEMS.00115-19-F0001.JPEG

Kleiner, M., Thorson, E., Sharp, C. E., Dong, X., Liu, D., Li, C., & Strous, M. (2017). Assessing species biomass contributions in microbial communities via metaproteomics. *Nature Communications 2017 8:1*, *8*(1), 1–14. https://doi.org/10.1038/S41467-017-01544-X

Lee, C. O., Boe-Hansen, R., Musovic, S., Smets, B., Albrechtsen, H. J., & Binning, P. (2014). Effects of dynamic operating conditions on nitrification in biological rapid sand filters for drinking water treatment. *Water Research*, *64*(m), 226–236. https://doi.org/10.1016/j.watres.2014.07.001

Li, D., Liu, C. M., Luo, R., Sadakane, K., & Lam, T. W. (2015). MEGAHIT: An ultra-fast single-node solution for large and complex metagenomics assembly via succinct de Bruijn graph. *Bioinformatics*, *31*(10), 1674–1676. https://doi.org/10.1093/bioinformatics/btv033

Li, H., Handsaker, B., Wysoker, A., Fennell, T., Ruan, J., Homer, N., Marth, G., Abecasis, G., & Durbin, R. (2009). The Sequence Alignment/Map format and SAMtools. *Bioinformatics*, *25*(16), 2078–2079. https://doi.org/10.1093/BIOINFORMATICS/BTP352

Lydmark, P., Lind, M., Sörensson, F., & Hermansson, M. (2006). Vertical distribution of nitrifying populations in bacterial biofilms from a full-scale nitrifying trickling filter. *Environmental Microbiology*, *8*(11), 2036–2049. https://doi.org/10.1111/j.1462-2920.2006.01085.x

Marcus, D. N., Pinto, A., Anantharaman, K., Ruberg, S. A., Kramer, E. L., Raskin, L., & Dick, G. J. (2017). Diverse manganese(II)-oxidizing bacteria are prevalent in drinking water systems. *Environmental Microbiology Reports*, *9*(2), 120–128. https://doi.org/10.1111/1758-2229.12508

McAllister, S. M., Polson, S. W., Butterfield, D. A., Glazer, B. T., Sylvan, J. B., & Chan, C. S. (2020). Validating the Cyc2 Neutrophilic Iron Oxidation Pathway Using Meta-omics of Zetaproteobacteria Iron Mats at Marine Hydrothermal Vents. *MSystems*, *5*(1). https://doi.org/10.1128/MSYSTEMS.00553-19

Nitzsche, K. S., Weigold, P., Lösekann-Behrens, T., Kappler, A., & Behrens, S. (2015). Microbial community composition of a household sand filter used for arsenic, iron, and manganese removal from groundwater in Vietnam. *Chemosphere*, *138*, 47–59. https://doi.org/10.1016/j.chemosphere.2015.05.032

Pacini, V. A., Ingallinella, A. M., & Sanguinetti, G. (2005). Removal of iron and manganese using biological roughing up flow filtration technology. *Water Research*, *39*(18), 4463–4475. https://doi.org/10.1016/j.watres.2005.08.027

Palomo, A., Jane Fowler, S., Gülay, A., Rasmussen, S., Sicheritz-Ponten, T., & Smets, B. F. (2016). Metagenomic analysis of rapid gravity sand filter microbial communities suggests novel physiology of Nitrospira spp. *ISME Journal*, *10*(11), 2569–2581. https://doi.org/10.1038/ismej.2016.63

Pinto, A. J., Marcus, D. N., Ijaz, U. Z., Bautista-de lose Santos, Q. M., Dick, G. J., & Raskin, L. (2016). Metagenomic Evidence for the Presence of Comammox Nitrospira -Like Bacteria in a Drinking Water System. *MSphere*, *1*(1). https://doi.org/10.1128/msphere.00054-15

Poghosyan, L., Koch, H., Frank, J., van Kessel, M. A. H. J., Cremers, G., van Alen, T., Jetten, M. S. M., Op den Camp, H. J. M., & Lücker, S. (2020). Metagenomic profiling of ammonia- and methane-oxidizing microorganisms in two sequential rapid sand filters. *Water Research*, *185*, 116288. https://doi.org/10.1016/j.watres.2020.116288

Purkhold, U., POMMERENING-ROSER, A., JURETSCHKO, S., SCHMID, M. C., & WAGNER, H.-P. K. M. (2000). Phylogeny of All Recognized Species of Ammonia Oxidizers Based on Comparative 16S rRNA and. *Society*, *66*(12), 5368–5382.

Qin, S., Ma, F., Huang, P., & Yang, J. (2009). Fe (II) and Mn (II) removal from drilled well water: A case study from a biological treatment unit in Harbin. *Desalination*, *245*(1–3), 183–193. https://doi.org/10.1016/j.desal.2008.04.048

Qin, Y. Y., Li, D. T., & Yang, H. (2007). Investigation of total bacterial and ammonia-oxidizing bacterial community composition in a full-scale aerated submerged biofilm reactor for drinking water pretreatment in China. *FEMS Microbiology Letters*, *268*(1), 126–134. https://doi.org/10.1111/j.1574-6968.2006.00571.x

Ramsay, L., Du, F., Lund, M., He, H., & Søborg, D. A. (2021). Grain displacement during backwash of drinking water filters. *Water Science and Technology: Water Supply*, *21*(1), 356–367. https://doi.org/10.2166/ws.2020.300

Romano, C. A., Zhou, M., Song, Y., Wysocki, V. H., Dohnalkova, A. C., Kovarik, L., Paša-Tolić, L., & Tebo, B. M. (2017). Biogenic manganese oxide nanoparticle formation by a multimeric multicopper oxidase Mnx. *Nature Communications*, *8*(1), 1–8. https://doi.org/10.1038/s41467-017-00896-8

Rost, B. (1999). Twilight zone of protein sequence alignments. *Protein Engineering, Design and Selection*, *12*(2), 85–94. https://doi.org/10.1093/PROTEIN/12.2.85

Schulze, W. X., Gleixner, G., Kaiser, K., Guggenberger, G., Mann, M., & Schulze, E. D. (2005). A proteomic fingerprint of dissolved organic carbon and of soil particles. *Oecologia*, *142*(3), 335–343. https://doi.org/10.1007/s00442-004-1698-9

Sharma, S. K., Petrusevski, B., & Schippers, J. C. (2005). Biological iron removal from groundwater: A review. *Journal of Water Supply: Research and Technology - AQUA*, *54*(4), 239–247. https://doi.org/10.2166/aqua.2005.0022

Stokke, R., Roalkvam, I., Lanzen, A., Haflidason, H., & Steen, I. H. (2012). Integrated metagenomic and metaproteomic analyses of an ANME-1-dominated community in marine cold seep sediments. *Environmental Microbiology*, *14*(5), 1333–1346. https://doi.org/10.1111/j.1462-2920.2012.02716.x

Tatari, K., Musovic, S., Gülay, A., Dechesne, A., Albrechtsen, H. J., & Smets, B. F. (2017). Density and distribution of nitrifying guilds in rapid sand filters for drinking water production: Dominance of Nitrospira spp. *Water Research*, *127*, 239–248. https://doi.org/10.1016/j.watres.2017.10.023

Tatari, K., Smets, B. F., & Albrechtsen, H. J. (2013). A novel bench-scale column assay to investigate site-specific nitrification biokinetics in biological rapid sand filters. *Water Research*, *47*(16), 6380–6387. https://doi.org/10.1016/j.watres.2013.08.005

Tatari, K., Smets, B. F., & Albrechtsen, H. J. (2016). Depth investigation of rapid sand filters for drinking water production reveals strong stratification in nitrification biokinetic behavior. *Water Research*, *101*, 402–410. https://doi.org/10.1016/j.watres.2016.04.073

Tekerlekopoulou, A. G., Pavlou, S., & Vayenas, D. V. (2013). Removal of ammonium, iron and manganese from potable water in biofiltration units: A review. *Journal of Chemical Technology and Biotechnology*, *88*(5), 751–773. https://doi.org/10.1002/jctb.4031

Uhl, W., & Gimbel, R. (2000). Dynamic modeling of ammonia removal at low temperatures in drinking water rapid filters. *Water Science and Technology*, *41*(4–5), 199–206. https://doi.org/10.2166/wst.2000.0445

Van Beek, C. G. E. M., Hiemstra, T., Hofs, B., Nederlof, M. M., Van Paassen, J. A. M., & Reijnen, G. K. (2012). Homogeneous, heterogeneous and biological oxidation of iron(II) in rapid sand filtration. *Journal of Water Supply: Research and Technology - AQUA*, *61*(1), 1–13. https://doi.org/10.2166/aqua.2012.033

Van Den Bossche, T., Kunath, B. J., Schallert, K., Schäpe, S. S., Abraham, P. E., Armengaud, J., Arntzen, M., Bassignani, A., Benndorf, D., Fuchs, S., Giannone, R. J., Griffin, T. J., Hagen, L. H., Halder, R., Henry, C., Hettich, R. L., Heyer, R., Jagtap, P., Jehmlich, N., … Muth, T. (2021). Critical Assessment of MetaProteome Investigation (CAMPI): a multi-laboratory comparison of established workflows. *Nature Communications*, *12*(1). https://doi.org/10.1038/S41467-021-27542-8

Vasimuddin, M., Misra, S., Li, H., & Aluru, S. (2019). Efficient architecture-aware acceleration of BWA-MEM for multicore systems. *Proceedings - 2019 IEEE 33rd International Parallel and Distributed Processing Symposium, IPDPS 2019*, 314–324. https://doi.org/10.1109/IPDPS.2019.00041

Von Meijenfeldt, F. A. B., Arkhipova, K., Cambuy, D. D., Coutinho, F. H., & Dutilh, B. E. (2019a). Robust taxonomic classification of uncharted microbial sequences and bins with CAT and BAT. *Genome Biology*, *20*(1), 1–14. https://doi.org/10.1186/S13059-019-1817-X/FIGURES/6

Von Meijenfeldt, F. A. B., Arkhipova, K., Cambuy, D. D., Coutinho, F. H., & Dutilh, B. E. (2019b). Robust taxonomic classification of uncharted microbial sequences and bins with CAT and BAT. *Genome Biology*, *20*(1). https://doi.org/10.1186/S13059-019-1817-X

Vries, D., Bertelkamp, C., Schoonenberg Kegel, F., Hofs, B., Dusseldorp, J., Bruins, J. H., de Vet, W., & van den Akker, B. (2017). Iron and manganese removal: Recent advances in modelling treatment efficiency by rapid sand filtration. *Water Research*, *109*, 35–45. https://doi.org/10.1016/j.watres.2016.11.032

W.W.J.M. de Vet. (2011). *Biological drinking water treatment of anaerobic groundwater in trickling filters* (Issue January 2011). TU Delft.

Wagner, F. B., Nielsen, P. B., Boe-Hansen, R., & Albrechtsen, H. J. (2016). Copper deficiency can limit nitrification in biological rapid sand filters for drinking water production. *Water Research*, *95*, 280–288. https://doi.org/10.1016/j.watres.2016.03.025

Wilmes, P., & Bond, P. L. (2004). The application of two-dimensional polyacrylamide gel electrophoresis and downstream analyses to a mixed community of prokaryotic microorganisms. *Environmental Microbiology*, *6*(9), 911–920. https://doi.org/10.1111/J.1462-2920.2004.00687.X

Wilmes, P., Heintz-Buschart, A., & Bond, P. L. (2015). A decade of metaproteomics: Where we stand and what the future holds. *Proteomics*, *15*(20), 3409–3417. https://doi.org/10.1002/pmic.201500183

Yang, H., Yan, Z., Du, X., Bai, L., Yu, H., Ding, A., Li, G., Liang, H., & Aminabhavi, T. M. (2020). Removal of manganese from groundwater in the ripened sand filtration: Biological oxidation versus chemical auto-catalytic oxidation. *Chemical Engineering Journal*, *382*, 123033. https://doi.org/10.1016/j.cej.2019.123033

Yang, L., Li, X., Chu, Z., Ren, Y., & Zhang, J. (2014). Distribution and genetic diversity of the microorganisms in the biofilter for the simultaneous removal of arsenic, iron and manganese from simulated groundwater. *Bioresource Technology*, *156*, 384–388. https://doi.org/10.1016/J.BIORTECH.2014.01.067
